# Cathelicidin-3 associated with serum extracellular vesicles enables early diagnosis of a transmissible cancer

**DOI:** 10.1101/2021.12.06.471373

**Authors:** Camila Espejo, Richard Wilson, Ruth J. Pye, Julian C. Ratcliffe, Manuel Ruiz-Aravena, Eduard Willms, Barrett W. Wolfe, Rodrigo Hamede, Andrew F. Hill, Menna E. Jones, Gregory M. Woods, A. Bruce Lyons

## Abstract

The identification of practical early diagnosis biomarkers is a cornerstone of improved prevention and treatment of cancers. Such a case is devil facial tumour disease (DFTD), a highly lethal transmissible cancer afflicting virtually an entire species, the Tasmanian devil (*Sarcophilus harrisii*). Despite a latent period that can exceed one year, to date DFTD diagnosis requires visual identification of tumour lesions. To enable earlier diagnosis, which is essential for the implementation of effective conservation strategies, we analysed the extracellular vesicle (EV) proteome of 87 Tasmanian devil serum samples. The antimicrobial peptide cathelicidin-3 (CATH3) was enriched in serum EVs of both devils with clinical DFTD (87.9% sensitivity and 94.1% specificity) and devils with latent infection (i.e., collected while overtly healthy, but 3-6 months before subsequent DFTD diagnosis; 93.8% sensitivity and 94.1% specificity). As antimicrobial peptides can play a variety of roles in the cancer process, our results suggest that the specific elevation of serum EV-associated CATH3 may be mechanistically involved in DFTD pathogenesis. This EV-based approach to biomarker discovery is directly applicable to improving understanding and diagnosis of a broad range of diseases in other species, and these findings directly enhance the capacity of conservation strategies to ensure the viability of the imperilled Tasmanian devil population.

## Introduction

Cancer is a condition that affects all multicellular species with differing degrees of susceptibility. One of the main challenges in oncology is a lack of diagnostic tools that allow for early detection of cancerous processes. Commonly, cancer diagnosis relies on biomarkers that are present in identified cancerous masses (solid biopsy), or present in the bodily fluids of the cancer patient (liquid biopsy). During the past decade, liquid biopsies have increasingly gained attention as a source of cancer biomarkers over traditional solid biopsies as they have increasing potential for early disease detection (Zhou et al., 2020). One approach increasingly used in liquid biopsies is the analysis of extracellular vesicles (EVs). EVs are nano-sized bilipid membrane structures that are released by all cells. EVs mediate intercellular communication, including mechanisms of cancer progression (Willms et al., 2018) via their functional cargo such as proteins, lipids, and nucleic acids (Maas et al., 2017).

EVs are a promising biomarker source as they are accessible from almost all bodily fluids (Colombo et al., 2014). They exhibit high sensitivity and specificity in cancer diagnosis and prognosis (Choi et al., 2015, Hoshino et al., 2020, Melo et al., 2015), and have organotrophic characteristics that may indicate organ-specific metastasis in bodily fluids (Hoshino et al., 2015). Further, EVs have stable biological activities as their cargo (which can mediate multiple cancer mechanisms) is protected from enzymatic degradation by a bilipid membrane (Boukouris and Mathivanan, 2015). Considering these advantages, researchers have expressed great enthusiasm in the molecular analysis of EVs as an approach to cancer biomarker discovery in liquid biopsies. Proteins are well-studied EV cargo (Zhou et al., 2020), as isolating EVs from serum can allow the enrichment and detection of a greater range of proteins that are otherwise masked by high-abundance serum/plasma proteins (Takov et al., 2019). Several EV protein biomarkers enabling early diagnosis of human cancer have been identified to date (Melo et al., 2015, Khan et al., 2012, Norouzi-Barough et al., 2020).

Commonly, cancer is understood as an individual disease, as tumours usually emerge and die with their hosts. However, there are several examples of transmissible cancers that have developed the capacity for tumour cells to be transmitted from one individual to another one as allografts (Ostrander et al., 2016). Like other infectious diseases, transmissible cancers become a health problem at the population level, even to the point of threatening populations with extinction. One such case is the devil facial tumour disease (DFTD) that affects the Tasmanian devil (*Sarcophilus harrisii*; herein ‘devil’). Since the first identification of DFTD in 1996, the disease has spread across more than 90% of the devils’ range, leading to an 82% decline in local densities and reducing the total population to as few as 16,900 individuals (Cunningham et al., 2021). Due to the high mortality and epidemic nature of DFTD, the Tasmanian devil was listed as endangered by the International Union for the Conservation of Nature in 2008 and is protected by both Tasmanian State and Australian Federal legislation (Hawkins et al., 2008). The cause of DFTD is a clonal cancer of Schwann cell origin that is transmitted as a malignant tissue transplant among devils through bites (Hamede et al., 2013, Murchison et al., 2010). DFTD is a lethal cancer, almost always killing its host within 6 to 12 months after clinical presentation of tumours on facial, oral and neck regions (Hamede et al., 2012). A second transmissible cancer (DFT2), also of Schwann cell origin, was reported in 2016 (Pye et al., 2016). In this manuscript, DFTD refers to the transmissible cancer identified in 1996.

DFTD is currently diagnosed by the appearance of macroscopic tumours and subsequent confirmation in the laboratory on the basis of positive staining for periaxin, karyotype aberrance, and PCR of tumour biopsies (Tovar et al., 2011, Kwon et al., 2018). However, there is direct evidence that DFTD has a long latent period as devils can develop tumours between 3 to 13 months after initial exposure to the disease (Save the Tasmanian Devil Program, 2017). McCallum et al. (McCallum et al., 2009) suggested that the disease is unlikely to spread between individuals prior to the development of clinical signs, however this assumption has not been validated due to the lack of a preclinical test. In an effort to identify DFTD serum biomarkers that could potentially serve to predict preclinical stages, Kau et al. (Karu et al., 2016) demonstrated that a panel of fibrinogen peptides and seven metabolites could differentiate devils with overt DFTD from healthy controls with high sensitivity and specificity. Another study found elevated levels of the receptor tyrosine-protein kinase ERBB3 in the serum of devils infected with DFTD compared to healthy controls (Hayes et al., 2017). Despite the potential value of serum biomarkers for DFTD diagnosis, neither study confirmed their findings in samples from latent DFTD devils (3 to 13 months prior to clinical manifestation of tumours). The discovery and validation of a biomarker for early DFTD diagnosis would greatly improve the capacity for DFTD surveillance and population management and could ultimately assist in recovering devil numbers in wild populations.

To enable the preclinical diagnosis of DFTD, in this study we analysed the proteome of EVs derived from the serum of devils collected over five years of quarterly devil trapping expeditions at several remote field sites in Tasmania. The longitudinal nature of this long-term monitoring program allowed the collection of serum from devils during the presumed “latent period”, i.e., samples collected while devils were clinically healthy (no palpable tumour masses), 3–6 months prior to subsequent recapture and clinical diagnosis of DFTD. We included EV samples from three classes of wild devils: those with clinically diagnosed overt DFTD, these devils in presumed latent stage of DFTD infection, herein: “latent”), and healthy devils from an offshore island population isolated from DFTD. Captive devils never exposed to DFTD were also included as healthy controls. These samples were divided into discovery and validation cohorts for the identification of DFTD biomarkers that would enable early detection with serum collected during routine.

## Results

### Characterisation of EVs derived from Tasmanian devil serum

First, we used size exclusion chromatography columns to isolate extracellular vesicles from serum samples of healthy (DFTD free controls) and DFTD infected devils in different stages of the disease (Table 1). Transmission electron microscopy (TEM) and nanoparticle tracking analysis (NTA) were used to evaluate the morphology and size of isolated extracellular vesicles. TEM images confirmed the presence of EV structures in all disease stages and healthy controls, showing a typical EV morphology as closed vesicles with a cup-shaped structure as described in other studies (Figure 1A and Supplementary figure 1; Rikkert et al., 2019). NTA demonstrated the presence of a heterogeneous nanoparticle population with a small to medium size distribution, which did not differ based on DFTD clinical stage (Figure 1B). The different clinical stages of the disease were classified according to tumour volumes (see “methods”). Although health status/DFTD stage had a significant effect on the total number of nanoparticles (one-way ANOVA p = 0.02), no significant pairwise differences between groups were found (Figure 1C).

**Table 1.** Summary of Tasmanian devil cohorts used in this study.

|  | Discovery cohort |  |  | Validation cohort |  |  |  |  |  | Total Devils |
| --- | --- | --- | --- | --- | --- | --- | --- | --- | --- | --- |
|  | All | Control | Adv. DFT1 | All | Control | Latent DFT1 | Early DFT1 | Med. DFT1 | Late DFT1 |  |
| **Total Cases** | 22 | 10 | 12 | 65 | 17 | 15 | 17 | 14 | 2 | 87 |
| **Age** |  |  |  |  |  |  |  |  |  |  |
| Adult | 22 | 10 | 12 | 40 | 8 | 6 | 12 | 13 | 1 | 62 |
| Juvenile | 0 | 0 | 0 | 25 | 9 | 9 | 5 | 1 | 1 | 25 |
| **Sex** |  |  |  |  |  |  |  |  |  |  |
| Male | 11 | 5 | 6 | 34 | 10 | 8 | 8 | 8 | 0 | 45 |
| Female | 11 | 5 | 6 | 31 | 7 | 7 | 9 | 6 | 2 | 42 |
| **Location** |  |  |  |  |  |  |  |  |  |  |
| West Takone | 9 | 0 | 9 | 38 | 0 | 11 | 15 | 10 | 2 | 47 |
| Wilmot | 3 | 0 | 3 | 10 | 0 | 4 | 2 | 4 | 0 | 13 |
| Maria Island | 0 | 0 | 0 | 17 | 17 | 0 | 0 | 0 | 0 | 17 |
| Richmond* | 6 | 6 | 0 | 0 | 0 | 0 | 0 | 0 | 0 | 6 |
| Bonorong* | 4 | 4 | 0 | 0 | 0 | 0 | 0 | 0 | 0 | 4 |
| **Tumour volume (ml)** |  |  |  |  |  |  |  |  |  |  |
| Median | NA | NA | 24.00 | NA | NA | NA | 0.76 | 23.30 | 41.15 | NA |
| IQR | NA | NA | 55.82 - 67.60 | NA | NA | NA | 0.58 - 1.35 | 14.70 - 30.29 | 33.60 - 48.69 | NA |
| **Corporal condition** |  |  |  |  |  |  |  |  |  |  |
| Emaciated | 0 | 0 | 1 | 0 | 0 | 0 | 0 | 0 | 1 | 2 |
| Moderately thin | 0 | 0 | 6 | 0 | 0 | 0 | 0 | 2 | 0 | 8 |
| Average | 0 | 0 | 5 | 0 | 17 | 14 | 13 | 12 | 1 | 62 |
| Good | 0 | 10 | 0 | 0 | 0 | 1 | 4 | 0 | 0 | 15 |
| Obese | 0 | 0 | 0 | 0 | 0 | 0 | 0 | 0 | 0 | 0 |

**Control:** healthy devils never exposed to DFTD. **IQR:** interquartile range.
\*Captive holding facilities.

**Fig. 1.**
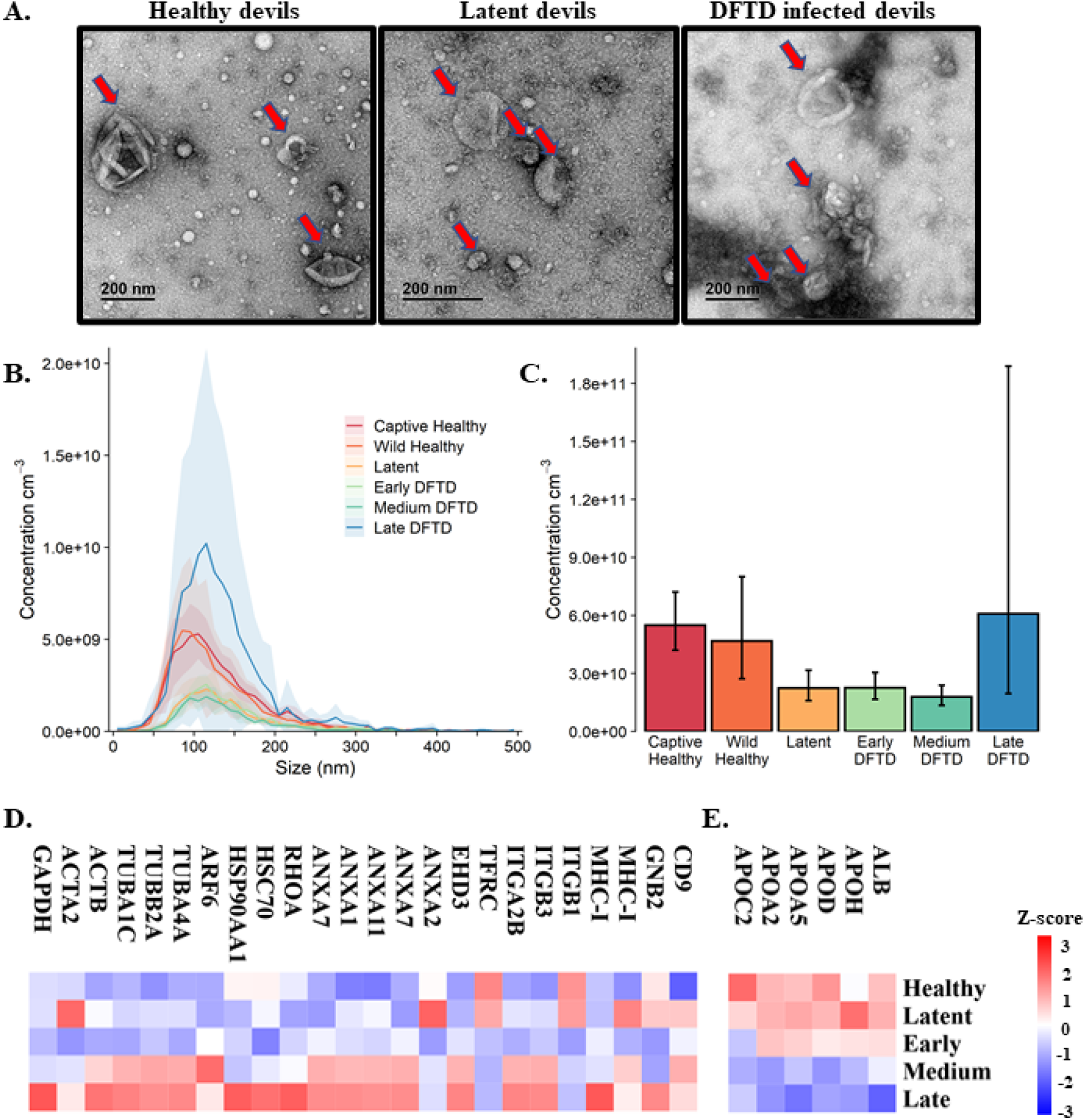
Characterisation of extracellular vesicles (EVs) derived from Tasmanian devil serum. **A**. Transmission electron microscopy images for EVs isolated from serum of healthy control wild devils from an isolated disease-free population (n=4), latent DFTD devils (n=4) and DFTD infected devils (n=4). Red arrows indicate EV structures. **B**. Size distribution profiles determined by nanoparticle tracking analysis (NTA) of EVs isolated from serum of captive (n=4) and wild (n=4) healthy control devils, latent DFTD devils (n=4), and DFTD infected devils in early (n=4), medium (n=4) and late stages (n=4). Shaded areas represent 95% confidence intervals. **C**. EV concentrations of the same NTA groups. Error bars represent 95% confidence intervals. **D**. Heat map of intensity values of commonly recovered EV proteins, and **E**. serum contaminants (albumin and five lipoproteins) found in EV samples derived from healthy controls (captive and wild) devils (n=27), latent DFTD devils (n=15), and devils in early (n=17), medium (n=15), and late (n=13) stages of DFTD.

A proteome dataset comprising combined discovery and validation cohorts (n=87) from the biomarker discovery process was generated by data independent acquisition mass spectrometry to gain an overview of the serum EV proteome and evaluate the presence of commonly recovered EV protein markers and serum contaminants (Thery et al., 2018). Of a total of 345 filtered proteins, 24 established EV markers were identified, including CD9, annexins, heat shock and major histocompatibility complex proteins (Kowal et al., 2016b) (Figure 1D). Serum-derived contaminants, which included albumin and five lipoproteins, all decreased in abundance as DFTD progressed (Figure 1E).

### Discovery of EV biomarkers for DFTD

For the biomarker discovery process, we first analysed the proteome of extracellular vesicles isolated from a cohort of serum samples from 22 devils (12 devils with advanced-stage DFTD; and 10 captive healthy controls) to identify EV protein DFTD biomarker candidates (Figure 2A and Table 1). Based on Student’s t-tests, 96 proteins (FDR corrected p < 0.05) were upregulated in EVs derived from DFTD infected devils relative to those from healthy controls (Figure 2B and Supplementary table 1A). Of these upregulated proteins, receiver operating characteristic (ROC) curve analysis identified 31 proteins with high accuracy (area under the ROC curve ≥ 0.9; Safari et al., 2016) to distinguish diseased from healthy individuals (Supplementary table 2). Proteins such as cathelicidin-3 (CATH3), connective tissue growth factor (CTGF) and complement component 5 (C5) were perfect classifiers of advanced-stage DFTD infected devils when compared to healthy controls (area under the ROC curve = 1, sensitivity and specificity = 100%; Figure 2C and Supplementary table 2). CATH3 was the most significantly upregulated protein in EVs derived from DFTD infected devils relative to healthy controls (p < 10e-6) with a 4.7-fold increase (Figures 2B and 2D). CTGF and C5 were significantly upregulated by 5.7- and 2.4-fold, respectively, in the DFTD infected devils compared to healthy controls (Figures 2B and 2D).

**Fig. 2.**
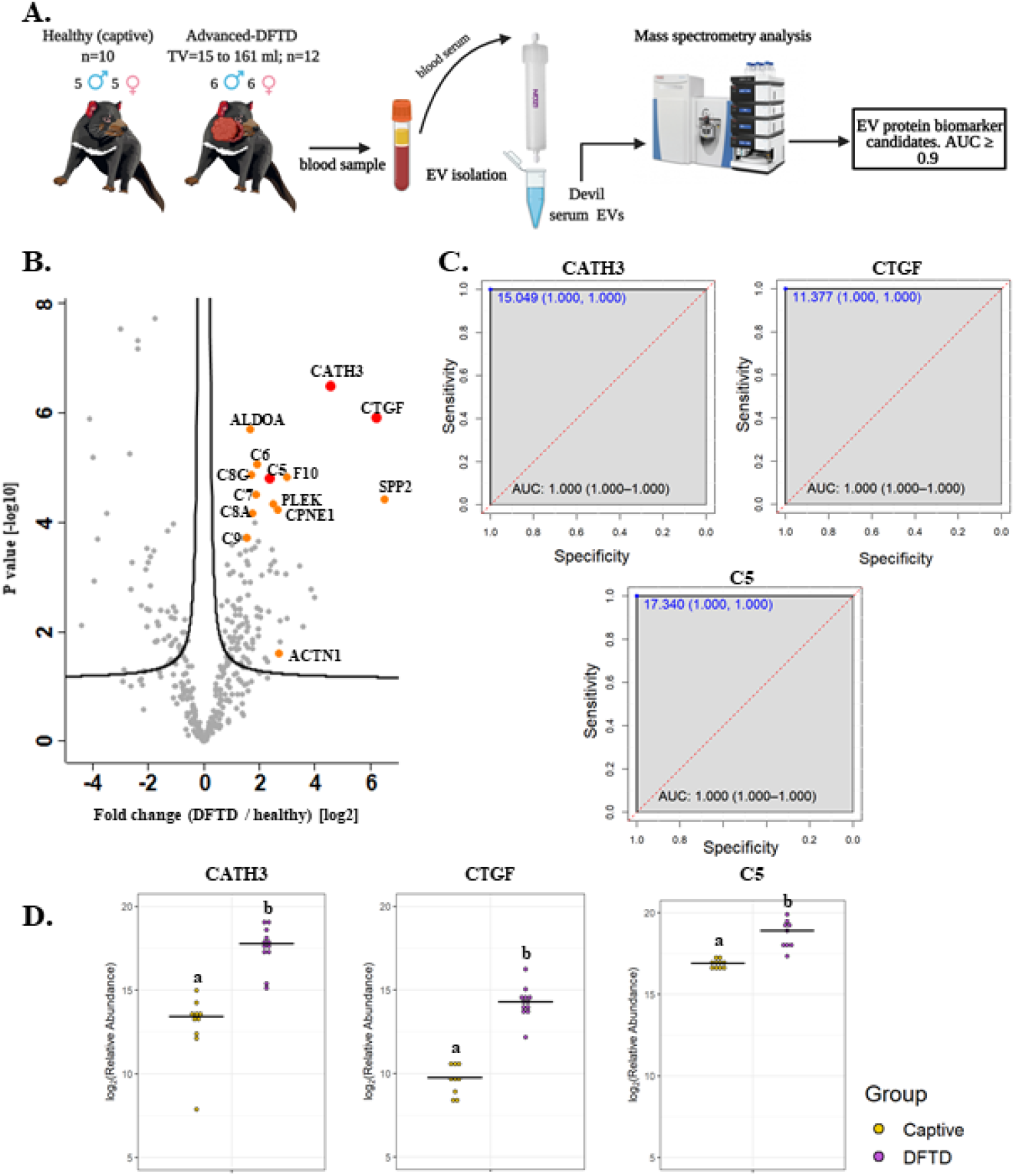
Discovery cohort. **A**. Extracellular vesicles (EVs) were isolated from 22 devil serum samples (12 DFTD-advanced and 10 healthy captive devils) by size exclusion chromatography columns. EVs were analysed by mass spectrometry to explore DFTD EV biomarker candidates. TV = tumour volume. **B**. Volcano plot of protein relative abundance fold changes (log_2_) between EVs derived from advanced stage DFTD and healthy devils vs fold change significance. Proteins denoted in red are perfect disease status classifiers (area under the receiver operating characteristic curve [AUC] = 1.0). Orange denotes proteins with an AUC > 0.95. **C**. Receiver operating characteristic curve analysis for the EV proteins CATH3, CTGF, and C5 (DFTD relative to healthy controls). The dashed red identity line indicates random performance. The cut-off values were determined using Youden’s index and are indicated in blue at the left top corner of the ROC curve and specificity and sensitivity are indicated in brackets, respectively. **D**. Dot plots depicting the relative abundance of the proteins CATH3, CTGF and C5 obtained from healthy animals and devils with advanced stages of DFTD, different letters “a” and “b” indicate significant differences between groups (Student’s t test, FDR-corrected p < 0.05).

To evaluate whether the upregulated proteins present in serum of advanced DFTD-infected devils relative to healthy controls were potentially released by DFTD cells, we used a proteomic database of EVs derived from cultured DFTD cells (Espejo et al., 2021). We found that of the 96 upregulated EV proteins derived from serum of DFTD infected devils relative to healthy controls, 19 of them overlapped with the proteins of EVs derived from DFTD cells that were upregulated relative to EVs derived from healthy fibroblasts (Supplementary figure 2). Six of these 19 proteins found in both cell culture EVs as well as serum proteomic databases yielded an area under the ROC curve greater than 0.9: F-actin-capping protein subunit alpha (CAPZA) and beta (CAPZB), profilin-1 (PFN1), fructose-bisphosphate aldolase A (ALDOA), tyrosine 3-monooxygenase/tryptophan 5-monooxygenase activation protein zeta (YWHAZ), and ARP3 actin related protein 3 (ACTR3) (Supplementary table 2 and Supplementary figure 2). However, none of the three perfect classifiers detected in the discovery cohort (CATH3, CTGF, and C5) were present in the EV cell culture database, suggesting an origin other than DFTD tumours.

### CATH3 and PFN1 as EV protein biomarkers for DFTD

To validate the discovery cohort results, the analysis of the proteome of EVs was repeated with an independent cohort of 33 DFTD-infected devils in different stages of the disease to test whether our potential EV protein biomarkers can identify animals in a broader range of cancer progression (Figure 3A and Table 1). We also included 17 healthy devils from a DFTD-free wild insurance population located on Maria Island as negative controls (Figure 3A and Table 1). Based on Student’s t-test analyses, 51 proteins (FDR-corrected p < 0.05) were upregulated in EVs derived from DFTD infected devils relative to healthy controls (Figure 3B and Supplementary table 1B). Of these 51 upregulated proteins, only four yielded an area under the ROC curve greater than 0.9 (Supplementary table 3). In agreement with the discovery cohort results, CATH3 and PFN1 were significantly upregulated in different stages of DFTD-infected devils relative to the wild healthy controls by 2.9- and 4.1-fold, respectively (Figures 3B and 3C). ROC curves indicated that CATH3 and PFN1 classified devils with DFTD with 87.9% and 90.9% sensitivity and 94.1% and 88.2% specificity, respectively (Figure 3D and Supplementary table 3). Unlike CATH3, PFN1 was detected in the cell culture DFTD EV database (Supplementary figure 2), suggesting a possible tumour origin.

**Fig. 3.**
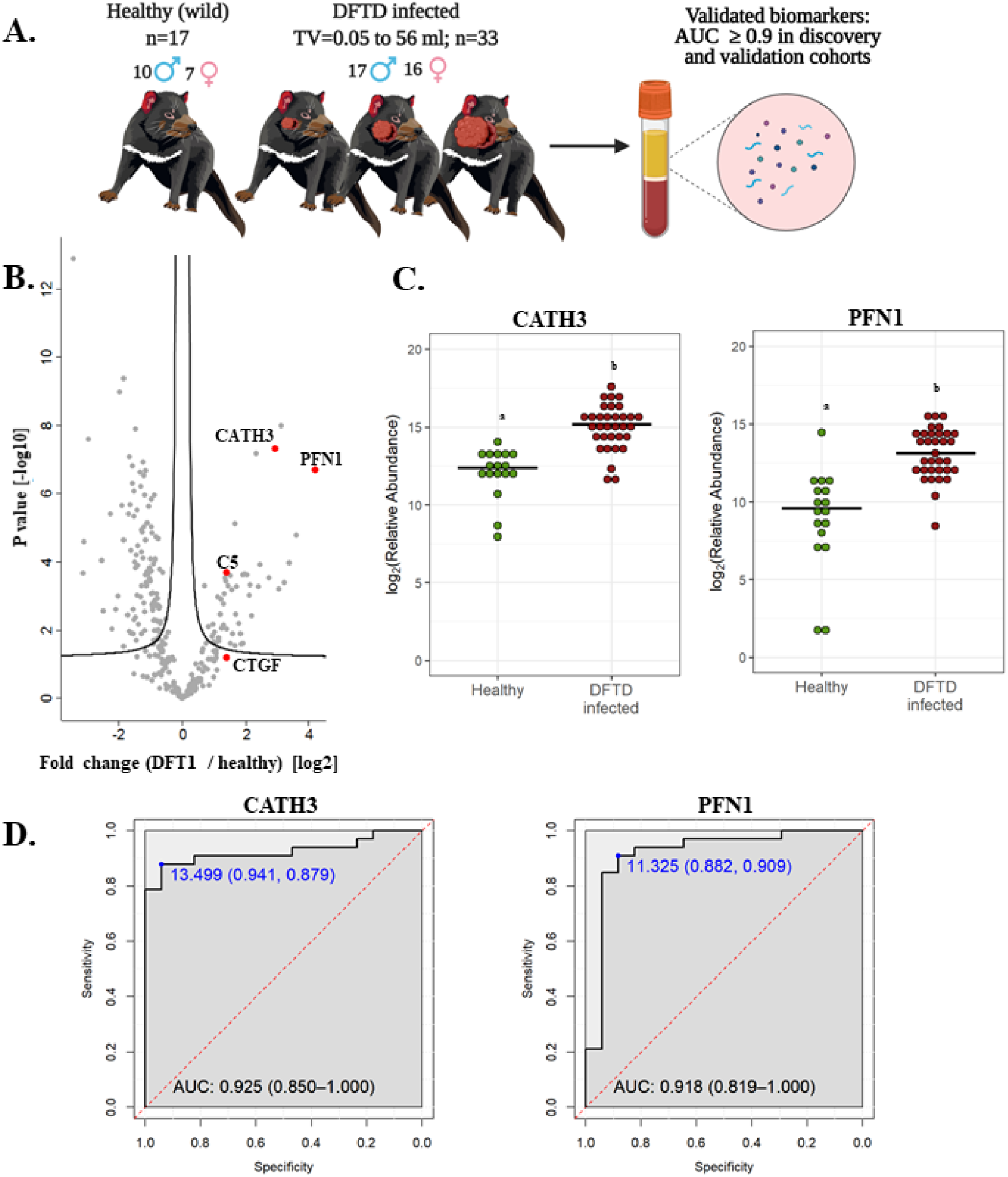
CATH3 and PFN1 EVs as biomarkers for DFTD. **A**. Isolated extracellular vesicles from serum samples of 50 devils (33 DFTD-infected and 17 healthy wild devils) were analysed by mass spectrometry to validate potential EV protein biomarkers detected in the discovery cohort results; TV= tumour volume. **B**. Volcano plot of protein relative abundance fold changes (log_2_) between EVs derived from serum of devils with different stages of DFTD (n=33) and healthy wild controls (n=17) vs fold change significance. **C**. Dot plot showing the relative abundance of CATH3 and PFN1 in the serum EVs of 17 wild healthy devils and 33 DFTD infected devils, different letters “a” and “b” indicate significant differences between groups (Student’s t test, FDR-corrected p < 0.05). **D**. Receiver operating characteristic curve analysis for CATH3 and PFN1 EVs (33 DFTD-infected animals vs 17 healthy controls). The dashed red line indicates random performance. The cut-off values were determined using Youden’s index and are indicated in blue at the left top corner of the ROC curve, and specificity and sensitivity are indicated in brackets, respectively.

In contrast, other protein candidates identified in the discovery cohort such as CTGF and C5 showed a reduced performance in distinguishing different stages of DFTD from the wild healthy controls, with a sensitivity of 48.5% and specificity of 88.2% for CTGF and 84.8% sensitivity and 70.6% specificity for C5 (Supplementary figure 3).

### CATH3 EVs detect latent stage DFTD 3 – 6 months before overt disease

Further analysis of EVs derived from serum samples of the validation cohort revealed that the levels of CATH3 in EV samples could successfully distinguish devils in latent stages of DFTD (n=15) from healthy wild individuals (n=17). Devils were presumed to be in the latent stage of DFTD as samples were collected 3 to 6 months before subsequent DFTD pathological and clinical diagnosis (Figure 4A and Table 1). Specifically, the levels of CATH3 were consistently upregulated in latent DFTD samples relative to the wild healthy group, following the same pattern revealed by the discovery and validation cohort results (Figures 4B and 4C). In contrast, PFN1 was not significantly upregulated in latent devils relative to healthy controls (Figures 4B and 4C).

**Fig. 4.**
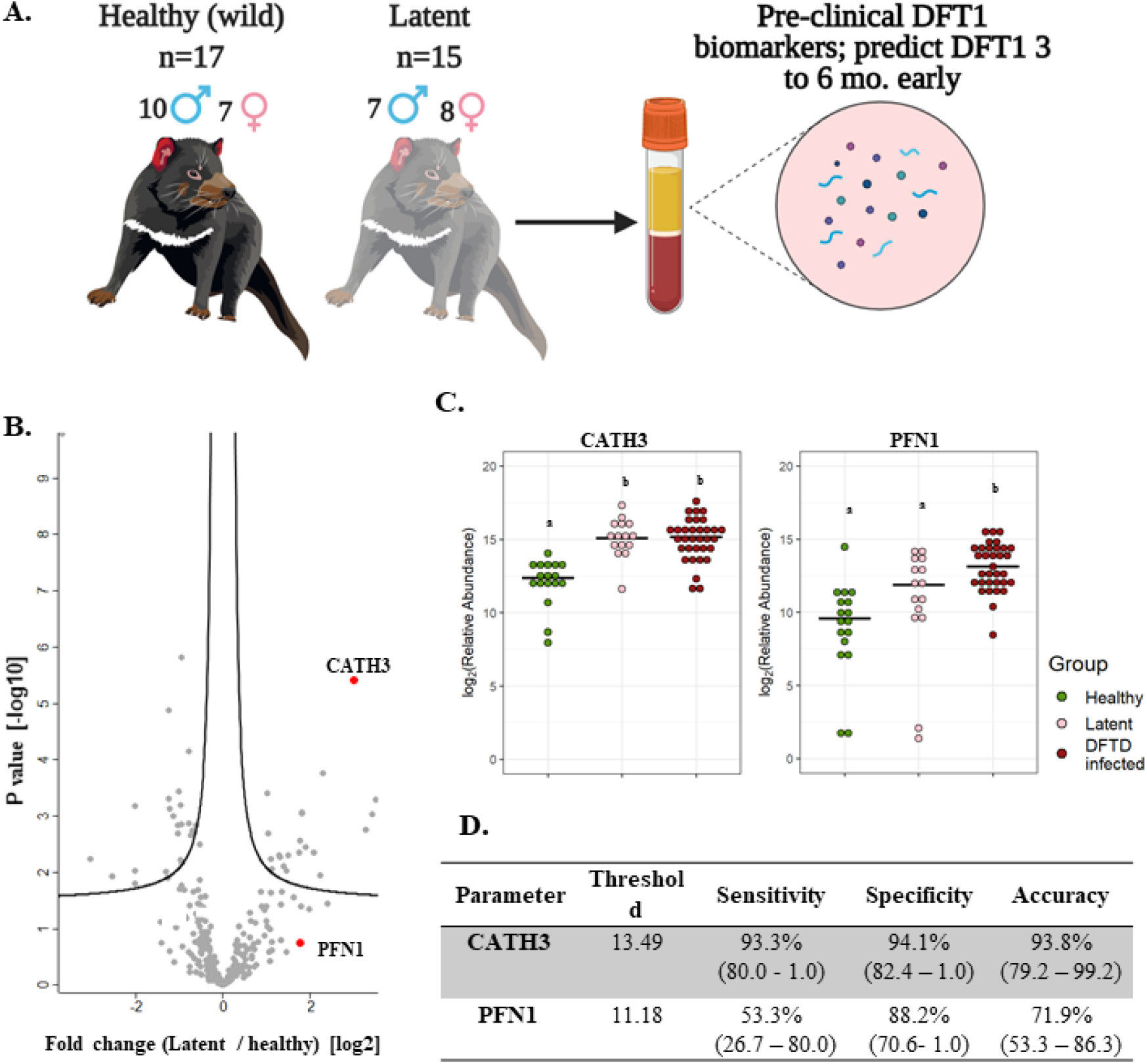
CATH3 EVs detect DFTD in latent stages. **A**. Isolated extracellular vesicles (EVs) from serum samples of 32 devils (15 DFTD latent and 17 healthy wild devils) were analysed to investigate whether the validated DFTD biomarkers can also serve to predict latent stage DFTD 3 to 6 months before overt DFTD. **B**. Volcano plot of protein relative abundance fold changes (log_2_) between EVs derived from serum of DFTD latent devils (n=15) and healthy wild controls (n=17) vs fold change significance. **C**. Dot plot showing the relative abundance of EV CATH3 and PFN1 detected in 17 wilds healthy, 15 latent, and 33 DFTD-infected devils, different letters “a” and “b” indicate significant pairwise differences between groups (i.e., groups denoted with the same letter are not significantly different; one-way ANOVA and Tukey post-hoc test, p < 0.05). **D**. Receiver operating curve analysis performed to classify latent devils (n=15) from healthy controls (n=17). Sensitivity and specificity were calculated (95% confidence intervals) for latent devils based on the protein threshold trained with the full validation dataset (n=50).

We calculated sensitivity, specificity, and accuracy of CATH3 and PFN1 to classify latent stages from healthy controls, using protein abundance cut-off values trained to distinguish DFTD infected devils from healthy controls calculated in the validation cohort. CATH3 exhibited a sensitivity of 93.3% and a specificity of 94.1% with an accuracy of 93.8% to differentiate latent stages from healthy controls, supporting its utility as a biomarker for all stages of DFTD and its potential use for early detection of this transmissible cancer (Figure 4D). In comparison with CATH3, the protein PFN1 was less effective in distinguishing devils in latent stages from healthy controls (Figure 4D).

### MYH10, TGFBI, and CTGF are associated with tumour burden

We used the filtered proteome dataset comprising combined discovery and validation cohorts to evaluate relationship between EV protein abundance and tumour volume. CTFG and C5 were significantly and positively correlated with tumour volume in DFTD infected devils (Figure 5A), which is consistent with their high predictive power to classify advanced-DFTD stages (large tumour volumes) from healthy individuals in the discovery biomarker phase. Myosin heavy chain 10 (MYH10), transforming growth factor beta induced (TGFBI) and CTGF were the proteins that correlated best with tumour volume (Figure 5A and Supplementary table 4), and their expression levels enhanced as tumour volume increases (Figure 5B). CATH3 and PFN1 did not demonstrate a significant positive correlation with tumour volume (Figure 5A) but showed a binary relationship with disease/healthy demonstrated in the discovery and validation cohorts.

**Fig. 5.**
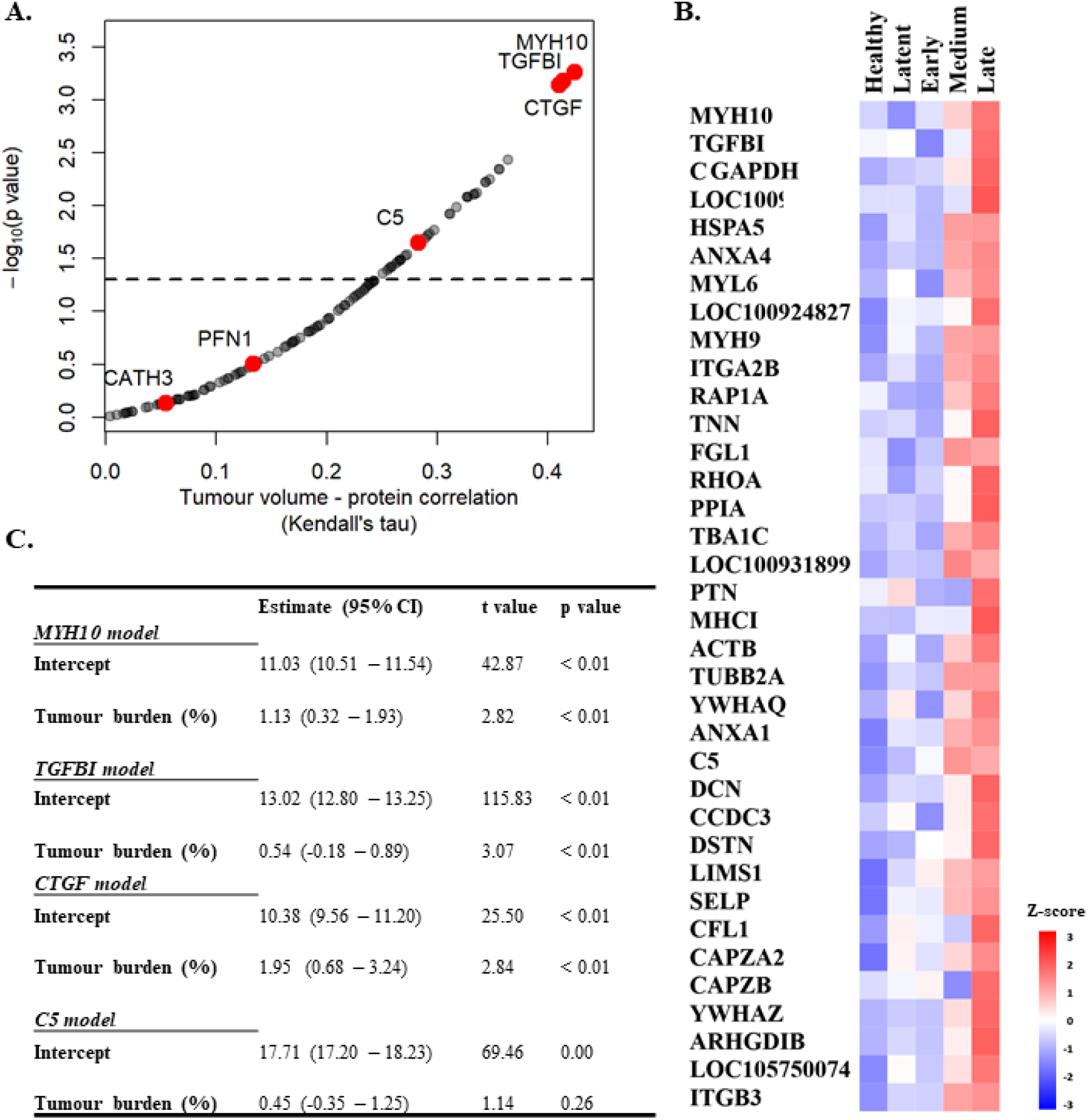
MYH10, TGFBI, and CTGF are associated with tumor burden. **A**. Kendall Correlation of proteins with tumor volumes (DFTD infected animals from the discovery and validation dataset, n=45). Proteins with a corrected p value <0.05 are plotted above the dashed and black line. **B**. Heat map representing EV proteins with significant Kendall correlations. Z-scored abundances were calculated from the mean of the relative abundance of each protein in each category (27 healthy (wild and captive), 15 latent, 17 early, 15 medium, and 13 late-DFTD devils). **C**. Linear regression models of % tumor burden as a predictor of MYH10, TGFBI, CTGF and C5 relative abundances.

Linear regressions were performed to evaluate the ability of tumour burden (as % of body mass) to predict MYH10, TGBI, CTGF, and C5 relative abundance values. Percent tumour burden was a significant predictor of CTGF (F (1,43) = 9.55, p < 0.01), TGFBI (F (1,43) = 9.41, p < 0.01), and MYH10 abundance (F (1,43) = 7.96, p < 0.01). Percent tumour burden explained a modest amount of variation in abundance of both CTGF and TGFBI (R^2^ = 0.18) and slightly less for MYH10 (R^2^ = 0.16). The models estimate CTGF, TGFBI, and MYH10 abundances enhance 1.95, 0.54- and 1.13-fold, respectively, for each 1% increase in tumour burden (Figure 5C).

## Discussion

The ongoing transmission of DFTD and the consequent decline of the Tasmanian devil population has been intensively investigated for the past 25 years. However, the sole method of diagnosis of this transmissible cancer still relies on the visual identification of tumours and confirmatory biopsy, despite previous efforts to develop preclinical diagnostic tests. Here, with contemporary methodology for isolation of extracellular vesicles and quantitative proteomics, we identified promising biomarker candidates from liquid biopsies with potential to predict the presence of this transmissible cancer at a preclinical stage. Specifically, the elevated expression of cathelicidin-3 (CATH3) in serum-derived EV samples of two independent cohorts had a high predictive power to detect DFTD. Further, CATH3 enrichment was detectable 3 to 6 months before tumours were visible or palpable, providing the first preclinical biomarker for DFTD and confirmation of a consistent latent period of DFTD infection. The preclinical detection of EV-associated CATH3 in routinely collected devil serum samples provides a means to improve the health management of endangered devils along with insights for the future development of mammalian cancer biomarkers.

Cathelicidins are a family of peptides with roles in antimicrobial responses (Zasloff, 2002). Relative to placental mammals, devils have a notable diversity of cathelicidin peptides, several of which are widely expressed in devil immune tissues, digestive, respiratory and reproductive tracts; milk and marsupium (i.e., pouch; Peel et al., 2016). Even though cathelicidins are thought to play important roles in the devil immune system, they have not been explored in DFTD pathogenesis. By contrast, the peptide LL-37, produced by the sole human cathelicidin gene, has been identified as a potential anti-tumour therapeutic agent for oral squamous cancer due to it causing apoptotic cell death, autophagy, and cell cycle arrest (Kuroda et al., 2015, Okumura et al., 2004). Conversely, other studies have suggested that LL-37 can promote cancer cell proliferation, migration, and tumour progression via activation of the MAPK/ERK signaling pathway (Weber et al., 2009, von Haussen et al., 2008). Interestingly, this pathway is interconnected with the ERRB-STAT3 axis, thought to be a primary mechanisms of tumorigenesis in DFTD (Kosack et al., 2019). These intriguing findings raise the possibility that CATH3 expression in the course of DFTD infection is associated with a protective response by the host animal’s innate immune system, or alternatively a yet undescribed evasion mechanism induced by the transmissible tumour.

As CATH3 was not identified in EVs derived from cultured DFTD tumour cells (Espejo et al., 2021), we propose that this early DFTD biomarker is likely associated with host cell derived EVs rather than those of tumour origin. EV protein cargo found in plasma/serum of cancer patients reflects the systemic effects of cancer, displaying markers not only associated with the primary tumour, but also the tumour microenvironment, distant organs, and the immune system (Hoshino et al., 2020). These EV protein signatures have also demonstrated diagnostic power in discriminating between healthy and cancer samples, indicating that host cell derived EVs can serve as sensitive cancer biomarkers. A host cell derived EV protein may have advantages for use as an early biomarker. Based on the finding that EV associated CATH3 abundance was independent of tumour volume, and the consistent upregulation of CATH3 across latent and overt DFTD stages relative to healthy samples, we propose that the increase in CATH3 arises from a uniform host response to this clonal cancer rather than of tumour cell origin. This independence of tumour volume is a desired feature for an early cancer biomarker as its sensitivity will be less dependent on a minimum tumour burden.

In contrast with the likely host origin of CATH3, we found that profilin-1 (PFN1), the overt DFTD biomarker found in this study, was highly expressed in EVs derived from DFTD cells *in vitro* (Espejo et al., 2021). As DFTD cells have a Schwann cell origin (Murchison et al., 2010), the upregulation of the actin-binding protein PFN1 is not surprising as it is required for Schwann cell development and migration (Montani et al., 2014). Additionally, PFN1 has been observed to be overexpressed in renal cell carcinoma (Minamida et al., 2011) and proposed as a urine biomarker for bladder cancer aggressiveness (Zoidakis et al., 2012). Considering these lines of evidence, we suggest that the upregulation of PFN1 in serum EVs isolated from overt DFTD devils likely originates from DFTD tumour cells. This is consistent with the poor performance of PFN1 to classify latent DFTD, considering tumour volume is presumably at a minimum at the preclinical disease stage.

Relative to the likely DFTD cell origin of PFN1 upregulation, a host origin of CATH3 may confer enhanced performance to classify preclinical DFTD but could also raise a concern regarding clinical specificity. Cathelicidins are also associated with inflammation and secondary infections (Zanetti, 2005), and altered abundance of other cathelicidins has been associated with purely inflammatory diseases such as bovine mastitis (Boehmer et al., 2008). However, we found no evidence for elevated levels of CATH3 in 16 of the 17 serum EV samples from the wild devils used for our healthy cohort despite the elevated values of other common inflammatory markers such as C-reactive protein, serum amyloid P-component, and several complement proteins (Eckersall and Bell, 2010, Karasu et al., 2018; see Supplementary figure 4). The high expression of these inflammatory markers found in wild healthy devils relative to captive healthy individuals is most likely due to the high prevalence of injuries resulting from intra-species biting, a common social behaviour among devils (Hamede et al., 2013) that results in wounds susceptible to microbial infections. Thus, the high specificity of CATH3, but not other cathelicidins or common inflammatory markers strongly implies that CATH3 is not associated with general inflammation. We suggest investigating the potential mechanism of action of CATH3 in the pathogenesis and progression of DFTD to identify this marker’s role.

Of the proteins that were found at greater abundance in devil EV samples at the advanced DFTD stages, many were among the subset of well-characterized EV markers, such as heat shock proteins, annexins, and integrins (Théry et al., 2018, Kowal et al., 2016a). This is consistent with previous reports that found a strong correlation of general EV markers with advanced cancer stages, indicating their potential prognostic value (Peinado et al., 2012, Hoshino et al., 2015). The three proteins myosin heavy chain 10 (MYH10), transforming growth factor beta induced (TGFBI), and connective growth factor (CTGF) with the strongest correlations with tumour volume are not generic EV markers and were not found in EVs derived from DFTD cells *in vitro* (Espejo et al., 2021). However, these proteins have all been documented to be associated with aggressiveness of tumour progression. For instance, MYH10 is overexpressed in glioma cells and implicated in cell migration and invasion (Senol et al., 2015), and also has a pro-tumorigenic effect in a murine lung cancer model (Kim et al., 2015). High expression of TGFBI predicts poor prognosis in patients with colorectal and ovarian cancer (Zhu et al., 2015, Karlan et al., 2014), while it also promotes breast cancer metastasis (Fico and Santamaria□Martínez, 2020). High levels of CTGF expression correlate positively with glioblastoma growth (Pan et al., 2002), invasive melanoma behaviour (Kubo et al., 1998), poor prognosis in oesophageal adenocarcinoma (Koliopanos et al., 2002), aggressive behaviour of pancreatic cancer cells (Wenger et al., 1999), and bone metastasis in breast cancer (Kang et al., 2003). Thus, the mechanisms that induce high levels of MYH10, TGFBI and CTGF expression with late stages of DFTD warrant further investigation to better understand the pathogenesis of DFTD.

The implementation of CATH3 as a diagnostic EV biomarker for latent DFTD will enhance the capabilities for management and conservation actions, which may aid the recovery of devils in wild populations, ensuring this species can fulfil its ecological niche in the future. Firstly, it will ensure that only healthy wild devils will be introduced into insurance populations, which will significantly reduce the cost of maintaining devils in quarantine prior to release, which is currently required for at least one year (Save the Tasmanian Devil Program, 2017). Secondly, it will greatly improve the capacity of ongoing monitoring programs that are critical for early warning and response and underpin research on the epidemiology and evolutionary dynamics of this disease system. Finally, early detection of DFTD will improve the implementation of any potential vaccination or other therapeutic intervention in the future (Flies et al., 2020). Further studies are required to determine whether CATH3 is elevated in devils in DFTD-latent periods longer than 3 to 6 months as the evidence suggests more than one year of latency, to determine how far pre-diagnosis CATH3 expression can distinguish latent devils from healthy controls.

The results herein demonstrate that DFTD is a valuable cancer model for comparative oncology to explore cancer biomarkers, as it represents a way to examine the effect of a single genetically identical cancer on the EV profile of numerous individual animals, allowing for a level of replication not possible in other systems. Identifying a devil cathelicidin as an early DFTD biomarker could provide insight into cancer responses more broadly and represent a possible target for the development of anticancer drugs, given human antimicrobial peptides have been proposed as novel cancer biomarkers and therapeutics agents (Kuroda et al., 2015, Deslouches and Di, 2017, Jin and Weinberg, 2019, Silva et al., 2018). Characterising CATH3 expression in response to a single cancer in a natural system could offer insight into host cancer adaptation strategies, as antimicrobial peptides have shown rapid evolutionary diversification within species with specific anti-pathogen activities (Lazzaro et al., 2020). Finally, this marker also widens the scope of human and animal cancer studies to include non-tumour derived cancer markers that result from altered physiology during tumour development.

## Material and Methods

### Serum samples

The two phases of this study comprised proteomic analysis of a discovery cohort and then a validation cohort (Table 1). The discovery phase aimed to identify potential EV protein biomarkers for DFTD using a cohort of 12 DFTD infected devils and 10 healthy controls. DFTD infected devils were considered in advanced stages (mid-late) of the disease based on large tumour volumes (15 ml to 161 ml). Tumour volumes were calculated by the ellipsoid formula described by Ruiz-Aravena et al. (Ruiz-Aravena et al., 2018), utilising measures of length, width, and depth of each DFTD tumour. DFTD infected devils often present more than one tumour on multiple locations of the body. Therefore, total tumour volume was calculated by summing the volume of each tumour present at the time of sampling. The second phase was designed to validate the first phase data in an independent cohort and further investigate the potential biomarkers in preclinical, presumed DFTD latent devils. The validation cohort was composed of 17 healthy controls, 15 latent (preclinical) DFTD-infected devils, and 33 confirmed DFTD-infected devils at different clinical stages of the disease. Of these, 17 devils were sub-classified as early stage (tumour volumes from 0.05 ml to 2.63 ml), 14 as medium stage (tumour volumes from 5.0 ml to 40.73), and 2 as late stage (tumour volumes from 26 ml to 56 ml). The animal with 26 ml of tumour was categorised as late instead of medium DFTD-stage as it was emaciated and had to be euthanized. The samples from presumed latent devils were collected 3 to 6 months prior to confirmed diagnosis of DFTD and are herein referred to as “latent” (Table 1).

The serum samples of the DFTD-infected devils used in both phases of the study were collected from two wild populations at the Northwest of Tasmania on 10-day field expeditions every 3 months between February 2015 and August 2019 (Table 1). The serum samples of the healthy cohort were obtained from captive devils held in Bonorong Wildlife Sanctuary and Richmond facilities (discovery cohort; samples collected between 2018-2019) and from wild devils from a DFTD-free insurance population (validation cohort; samples collected between 2014 and 2015) (Table 1). As DFTD-induced extinction was a genuine concern predicted by mathematical and epidemiological models (McCallum et al., 2009), government managers established a wild-DFTD population on an isolated island free from DFTD, located in Maria Island on Tasmania’s east coast (Thalmann et al., 2016). Blood was obtained from conscious (wild devils) or anesthetised devils (captive devils) by venepuncture from either jugular or marginal ear vein (between 0.3 – 1 mL) and transferred into empty or clot activating tubes. After a maximum of ∼five hours, samples were centrifuged at 1,000 *g* for 10 minutes and the serum was pipetted off and stored frozen at −20 C° (short term storage, up to 3 months) or −80 C° (long term storage, up to 6 years) until further use. All animal procedures were performed under a Standard Operating Procedure approved by the General Manager, Natural and Cultural Heritage Division, Tasmanian Government Department of Primary Industries, Parks, Water, and the Environment and under the auspices of the University of Tasmania Animal Ethics Committee (permit numbers A0017550, A0012513, A0013326, and A0015835).

### Extracellular vesicles purification

Serum samples were thawed on ice and 500 µl and 300 µl of serum were extracted for the discovery and validation cohort, respectively. The serum samples were firstly centrifuged at 1,500 *g* for 10 minutes at 4 °C to remove cells and debris. The samples were further centrifuged at 10,000 *g* for 10 minutes at 4 °C to pellet larger extracellular vesicles. The supernatant was taken and subjected immediately to size exclusion chromatography on qEV2 / 35nm columns (IZON) following the manufacturer’s instructions. Briefly, EVs were eluted in phosphate buffered saline (PBS) containing 0.05% sodium azide in eight fractions of 1 ml each after the collection of 14 ml of void volume and pooled. The EV samples were concentrated with Amicon Ultra-15 centrifugal filters (MWCO 100 kDa) to a final volume of 1 ml and stored in aliquots of 500 µl at −80 °C until future use.

### Transmission electron microscopy (TEM)

Copper TEM grids with a formvar-carbon support film (GSCU300CC-50, ProSciTech, Qld, Australia) were glow discharged for 60 seconds in an Emitech k950x with k350 attachment. Two 5 µl drops of EV suspension was pipetted onto each grid, allowed to adsorb for at least 30 seconds and blotted with filter paper after each. Two drops of 2% uranyl acetate were used to negatively stain the particles blotting after 10 seconds each time. Grids were then allowed to dry before imaging. Grids were imaged using a Joel JEM-2100 (JEOL (Australasia) Pty Ltd) transmission electron microscope equipped with a Gatan Orius SC 200 CCD camera (Scitek Australia).

### Nano particle tracking analysis (Zetaview)

EV size distribution and concentration were determined using a ZetaView PMX-120 nanoparticle analyzer (Particle Metrix, Inning am Ammersee, Germany) equipped with Zetaview Analyze Software version 8.05.12. Prior to measurement, the system was calibrated as per manufactures instruction with 100nm Nanospheres 3100A (Thermo Fisher Scientific). Measurements were performed in scatter mode, and for all measurements the cell temperature was maintained at 25 °C. Each sample was diluted in PBS to a final volume of 1 ml. Capture settings were sensitivity 80, shutter 100, and frame rate 30. Post-acquisition settings were minimum trace length 10, min brightness 30, min area 5, and max area 1000. Cell temperature was maintained at 25 °C for all measurements.

### Liquid chromatography and mass spectrometry analysis

#### Sample preparation

EV samples (500 µl aliquots) were thawed on ice and mixed with acetonitrile to a final concentration of 50% (v/v) and evaporated by a centrifugal vacuum concentrator to obtain EV proteins for mass spectrometry analysis. The EV protein samples were resuspended in 150 µl of denaturation buffer (7 M urea and 2 M thiourea in 40 mM Tris, pH 8.0).

Protein concentration was measured by EZQ protein quantification kit (Thermo Fisher Scientific), and 30 µg of protein from each sample was reduced with 10 mM dithiothreitol overnight at 4 °C. EV protein samples were alkylated the next day with 50 mM iodoacetamide for 2 hours at ambient temperature in the dark and then digested into peptides with 1.2 µg proteomics-grade trypsin/LysC (Promega) according to the SP3 protocol described by Hughes et al (2019). EV peptides were de-salted using ZipTips (Merck) according to the manufacturer’s directions.

#### High-pH peptide fractionation

A specific peptide spectral library was created for devil serum EVs using off-line high-pH fractionation. A pooled peptide sample (180 µg) composed of aliquots of each EV sample from the discovery cohort (n=22 individuals) was desalted with Pierce desalting spin columns (Thermo Fisher Scientific) according to manufacturer’s guidelines. The sample was evaporated to dryness and resuspended in 25 µl in HPLC loading buffer (2% acetonitrile with 0.05% TFA) and injected onto a 100 × 1 mm Hypersil GOLD (particle size 1.9 mm) HPLC column. Peptides were separated on an Ultimate 3000 RSLC system with micro fractionation and automated sample concatenation enabled at 30 µl/min with a 40 min linear gradient of 96% mobile phase A (water containing 1% triethylamine, adjusted to pH 9.6 utilizing acid acetic) to 50% mobile phase B (80% acetonitrile with 1% of triethylamine). The column was then washed in 90% buffer B and re-equilibrated in 96% buffer A for 8 minutes. Sixteen concatenated fractions were collected into 0.5 ml low-bind Eppendorf tubes, and then evaporated to dryness and reconstituted in 12 µl HPLC loading buffer.

#### Mass spectrometry – data-dependent acquisition (DDA)

Peptide fractions were analysed by nanoflow HPLC-MS/MS using an Ultimate 3000 nano RSLC system (Thermo Fisher Scientific) coupled with a Q-Exactive HF mass spectrometer fitted with a nano spray Flex ion source (Thermo Fisher Scientific) and controlled using Xcalibur software (version 4.3). Approximately 1 µg of each fraction was injected and separated using a 90-minute segmented gradient by preconcentration onto a 20 mm x 75 µm PepMap 100 C18 trapping column then separation on a 250 mm x 75 µm PepMap 100 C18 analytical column at a flow rate of 300 nL/min and held at 45°C. MS Tune software (version 2.9) parameters used for data acquisition were: 2.0 kV spray voltage, S-lens RF level of 60 and heated capillary set to 250 °C. MS1 spectra (390 - 1500 *m/z*) were acquired at a scan resolution of 120,000 followed by MS2 scans using a Top15 DDA method, with 30-second dynamic exclusion of fragmented peptides. MS2 spectra were acquired at a resolution of 15,000 using an AGC target of 2e5, maximum IT of 28ms and normalised collision energy of 30.

#### Mass spectrometry – data-independent acquisition (DIA)

Individual EV peptide samples were analysed by nanoflow HPLC-MS/MS using the instrumentation and LC gradient conditions described above but using DIA mode. MS1 spectra (390 - 1240 *m/z*) were acquired at 120 k resolution, followed by sequential MS2 scans across 26 DIA x 25 amu windows over the range of 397.5-1027.5 *m/z*, with 1 amu overlap between sequential windows. MS2 spectra were acquired at a resolution of 30,000 using an AGC target of 1e6, maximum IT of 55 ms and normalised collision energy of 27.

#### Proteomic database search

Both DDA-MS and DIA-MS raw files were processed using Spectronaut software (version 13.12, Biognosys AB). The specific library was generated using the Pulsar search engine to search DDA MS2 spectra against the *Sarcophilus harrisii* UniProt reference proteome (comprising 22,388 entries, last modified in August 2020). Spectral libraries were generated using all default software (BGS factory) settings, including N-terminal acetylation and methionine oxidation as variable modifications and cysteine carbamidomethylating as a fixed modification, up to two missed cleavages allowed and peptide, protein and PSM thresholds set to 0.01. For protein identification and relative quantitation between samples, DIA-MS data were processed according to BGS factory settings, with the exception that single-hit proteins were excluded. In the case of uncharacterized proteins, protein sequences provided by UniProt were blasted against the Tasmanian devil reference genome (GCA_902635505.1 mSarHar1.11) using the online NCBI protein blast tool (Wheeler et al., 2007).

### Statistical analysis

Spectronaut protein quantitation pivot reports, including protein description, gene names and UniProt accession numbers, were created for the discovery, validation, and combined datasets. The combined dataset includes the discovery and validation datasets and was used to search for EV protein markers suggested by the Minimal information for studies of extracellular vesicles 2018 (Thery et al., 2018), and to evaluate the relationship of EV proteins with tumour volume. The protein quantitation pivot reports were uploaded into Perseus software (version 1.6.10.50) for further data processing and statistical analysis. Quantitative values were log_2_ transformed and proteins filtered according to the number of valid values. The data was filtered in order that a valid value for a given protein was detected in ≥70% of samples in at least one group (i.e., discovery: control/diseased; validation: control/latent/early/advanced; combined: captive healthy/wild healthy/latent/early/medium/late). Remaining missing values were imputed with random intensity values for low-abundance proteins based on a normal abundance distribution using default Perseus settings. The filtered proteins in the discovery and validation datasets were considered for differential expression analyses of biomarker candidates, which was determined using two-tailed Student’s *t*-test with a permutation-based false discovery rate (FDR) controlled at 5% and s0 values set to 0.1 to exclude proteins with very small differences between means.

Significantly upregulated EV proteins from the filtered datasets were exported from Perseus and analysed using R 3.6.2 (R Core Team, 2019). The utility of each discovery dataset protein as a disease status classifier was investigated by subjecting healthy/disease cohort sample values of each to receiver operating characteristic (ROC) curve analysis to calculate their area under the curve, sensitivity, specificity, and accuracy with bootstrapped confidence intervals. The classification cut-off values were determined using Youden’s index. Discovery dataset proteins with a disease status classification area under the ROC curve greater than 0.9 were then investigated by ROC curve analyses, if present, in the validation dataset (excluding the latent samples). Proteins with ROC curves greater than 0.9 in the discovery and validation dataset were investigated in the latent cohort vs healthy wild controls using protein abundance cut-off values trained to distinguish DFTD infected devils from healthy controls calculated in the validation cohort. Kendall rank correlation was used to reveal significant correlations between protein abundance from the combined dataset and tumour volumes. Linear models were utilized to search for associations between the level of EV proteins and tumour burden, which was calculated by dividing total tumour mass by body weight minus total tumour mass and expressed in percentage. Tumour mass was calculated assuming a tumour density of 1.1 g per ml of volume as described by Ruiz-Aravena et al. (Ruiz-Aravena et al., 2018).

### Proteome of extracellular vesicles derived from DFTD cultured cells in vitro

To identify possible signals from DFTD tumors, an EV proteome database derived from cultured DFTD cells was used to identify proteins in serum EVs that may originate from DFTD cells (Espejo et al., 2021). Proteins upregulated in DFTD EVs relative to healthy fibroblast EVs and in EVs derived from serum samples of DFTD-infected devils relative to healthy controls obtained in the discovery cohort were compared.

## Acknowledgments

The authors would like to acknowledge all the members of the devil and wild immunology group for their advice and guidance. We would like to thank Ginny Ralph for providing care of captive devils, the Bonorong Wildlife Sanctuary for providing access to Tasmanian devils and Dr Alexandre Kreiss for collecting the blood and to the Save the Tasmanian Devil Program for provision of samples. We are grateful of Phil and Marita Crombie from Mistover Cottage in Yolla for providing accommodation and logistic support during fieldwork. We thank volunteers who helped with data collection. We also thank David Gell, Cherie Blenkiron and Kirsty Danielson for their comments in the manuscript.

## References

Boehmer, J., Bannerman, D., Shefcheck, K. & Ward, J. 2008. Proteomic analysis of differentially expressed proteins in bovine milk during experimentally induced Escherichia coli mastitis. Journal of dairy science, 91, 4206–4218.

Boukouris, S. & Mathivanan, S. 2015. Exosomes in bodily fluids are a highly stable resource of disease biomarkers. PROTEOMICS–Clinical Applications, 9, 358–367.

Choi, D. S., Kim, D. K., Kim, Y. K. & Gho, Y. S. 2015. Proteomics of extracellular vesicles: exosomes and ectosomes. Mass spectrometry reviews, 34, 474–490.

Colombo, M., Raposo, G. & Théry, C. 2014. Biogenesis, secretion, and intercellular interactions of exosomes and other extracellular vesicles. Annual review of cell and developmental biology, 30, 255–289.

Cunningham, C. X., Comte, S., Mccallum, H., Hamilton, D. G., Hamede, R., Storfer, A., Hollings, T., Ruiz-Aravena, M., Kerlin, D. H., Brook, B. W., Hocking, G. & Jones, M. E. 2021. Quantifying 25 years of disease-caused declines in Tasmanian devil populations: host density drives spatial pathogen spread. Ecol Lett, 24, 958–969.

Deslouches, B. & Di, Y. P. 2017. Antimicrobial peptides with selective antitumor mechanisms: prospect for anticancer applications. Oncotarget, 8, 46635.

Eckersall, P. & Bell, R. 2010. Acute phase proteins: Biomarkers of infection and inflammation in veterinary medicine. The veterinary journal, 185, 23–27.

Espejo, C., Wilson, R., Willms, E., Ruiz-Aravena, M., Pye, R. J., Jones, M. E., Hill, F., Woods, G. M. & Lyons, A. B. 2021. Extracellular vesicle proteomes of two transmissible cancers of Tasmanian devils reveal tenascin-C as a serum-based differential diagnostic biomarker. Cell Mol Life Sci.

Fico, F. & Santamaria□martínez, A. 2020. TGFBI modulates tumour hypoxia and promotes breast cancer metastasis. Molecular oncology, 14, 3198–3210.

Flies, A. S., Flies, E. J., Fox, S., Gilbert, A., Johnson, S. R., Liu, G.-S., Lyons, A. B., Patchett, A. L., Pemberton, D. & Pye, R. J. 2020. An oral bait vaccination approach for the Tasmanian devil facial tumor diseases. Expert review of vaccines, 19, 1–10.

Hamede, R., Lachish, S., Belov, K., Woods, G., Kreiss, A., Pearse, A. M., LAZENBY,, Jones, M. & Mccallum, H. 2012. Reduced Effect of Tasmanian Devil Facial Tumor Disease at the Disease Front. Conservation Biology, 26, 124–134.

Hamede, R. K., Mccallum, H. & Jones, M. 2013. Biting injuries and transmission of Tasmanian devil facial tumour disease. J Anim Ecol, 82, 182–90.

Hawkins, C., Mccallum, H., Mooney, N., Jones, M. & Holdsworth, M. 2008. Sarcophilus harrisii. The IUCN Red List of threatened species 2008: e. T40540A10331066.

Hayes, D. A., Kunde, D. A., Taylor, R. L., Pyecroft, S. B., Sohal, S. S. & Snow, E. T. 2017. ERBB3: A potential serum biomarker for early detection and therapeutic target for devil facial tumour 1 (DFT1). PloS one, 12, e0177919.

Hoshino, A., Costa-Silva, B., Shen, T. L., Rodrigues, G., Hashimoto, A., Tesic Mark, M., Molina, H., Kohsaka, S., Di Giannatale, A., Ceder, S., Singh, S., Williams, C., Soplop, N., Uryu, K., Pharmer, L., King, T., Bojmar, L., Davies, A. E., Ararso, Y., Zhang, T., Zhang, H., Hernandez, J., Weiss, J. M., Dumont-Cole, V. D., Kramer, K., Wexler, L. H., Narendran, A., Schwartz, G. K., Healey, J. H., Sandstrom, P., Labori, K. J., Kure, E. H., Grandgenett, P. M., Hollingsworth, M. A., De Sousa, M., Kaur, S., Jain, M., Mallya, K., Batra, S. K., Jarnagin, W. R., Brady, M. S., Fodstad, O., Muller, V., Pantel, K., Minn, A. J., Bissell, M. J., Garcia, B. A., Kang, Y., Rajasekhar, V. K., Ghajar, C. M., Matei, I., Peinado, H., Bromberg, J. & Lyden, D. 2015. Tumour exosome integrins determine organotropic metastasis. Nature, 527, 329–35.

Hoshino, A., Kim, H. S., Bojmar, L., Gyan, K. E., Cioffi, M., Hernandez, J., Zambirinis, C. P., Rodrigues, G., Molina, H., Heissel, S., Mark, M. T., Steiner, L., Benito-Martin, A., Lucotti, S., Di Giannatale, A., Offer, K., Nakajima, M., Williams, C., Nogues, L., Pelissier Vatter, F. A., Hashimoto, A., Davies, A. E., Freitas, D., Kenific, C. M., Ararso, Y., Buehring, W., Lauritzen, P., Ogitani, Y., Sugiura, K., Takahashi, N., Aleckovic, M., Bailey, K. A., Jolissant, J. S., Wang, H., Harris, A., Schaeffer, L. M., Garcia-Santos, G., Posner, Z., Balachandran, V. P., Khakoo, Y., Raju, G. P., Scherz, A., Sagi, I., Scherz-Shouval, R., Yarden, Y., Oren, M., Malladi, M., Petriccione, M., De Braganca, K. C., Donzelli, M., Fischer, C., Vitolano, S., Wright, G. P., Ganshaw, L., Marrano, M., Ahmed, A., Destefano, J., Danzer, E., Roehrl, M. H. A., Lacayo, N. J., Vincent, T. C., Weiser, M. R., Brady, M. S., Meyers, P. A., Wexler, L. H., Ambati, S. R., Chou, A. J., Slotkin, E. K., Modak, S., Roberts, S. S., Basu, E. M., Diolaiti, D., Krantz, B. A., Cardoso, F., Simpson, A. L., Berger, M., Rudin, C. M., Simeone, D. M., Jain, M., Ghajar, C. M., Batra, S. K., Stanger, B. Z., Bui, J., Brown, K. A., Rajasekhar, V. K., Healey, J. H., De Sousa, M., Kramer, K., Sheth, S., Baisch, J., Pascual, V., Heaton, T. E., La Quaglia, M. P., Pisapia, D. J., Schwartz, R., Zhang, H., Liu, Y., Shukla, A., Blavier, L., Declerck, Y. A., et al. 2020. Extracellular Vesicle and Particle Biomarkers Define Multiple Human Cancers. Cell, 182, 1044–1061 e18.

Hughes, C. S., Moggridge, S., Muller, T., Sorensen, P. H., Morin, G. B. & Krijgsveld, J. 2019. Single-pot, solid-phase-enhanced sample preparation for proteomics experiments. Nat Protoc, 14, 68–85.

Jin, G. & Weinberg, A. Human antimicrobial peptides and cancer. Seminars in cell & developmental biology, 2019. Elsevier, 156–162.

Kang, Y., Siegel, P. M., Shu, W., Drobnjak, M., Kakonen, S. M., Cordón-cardo, C., Guise, T. A. & Massagué, J. 2003. A multigenic program mediating breast cancer metastasis to bone. Cancer cell, 3, 537–549.

Karasu, E., Eisenhardt, S. U., Harant, J. & Huber-Lang, M. 2018. Extracellular vesicles: packages sent with complement. Frontiers in immunology, 9, 721.

Karlan, B. Y., Dering, J., Walsh, C., Orsulic, S., Lester, J., Anderson, L. A., Ginther, C. L., Fejzo, M. & Slamon, D. 2014. POSTN/TGFBI-associated stromal signature predicts poor prognosis in serous epithelial ovarian cancer. Gynecologic oncology, 132, 334–342.

Karu, N., Wilson, R., Hamede, R., Jones, M., Woods, G. M., Hilder, E. F. & Shellie, R. A. 2016. Discovery of biomarkers for Tasmanian Devil Cancer (DFTD) by metabolic profiling of serum. Journal of proteome research, 15, 3827–3840.

Khan, S., Jutzy, J. M., Valenzuela, M. M. A., Turay, D., Aspe, J. R., Ashok, A., Mirshahidi, S., Mercola, D., Lilly, M. B. & Wall, N. R. 2012. Plasma-derived exosomal survivin, a plausible biomarker for early detection of prostate cancer. PloS one, 7, e46737.

Kim, J. S., Kurie, J. M. & Ahn, Y.-H. 2015. BMP4 depletion by miR-200 inhibits tumorigenesis and metastasis of lung adenocarcinoma cells. Molecular cancer, 14, 1–11.

Koliopanos, A., Friess, H., Di Mola, F. F., Tang, W.-H., Kubulus, D., Brigstock, D., Zimmermann, A. & Büchler, M. W. 2002. Connective tissue growth factor gene expression alters tumor progression in esophageal cancer. World journal of surgery, 26, 420–427.

Kosack, L., Wingelhofer, B., Popa, A., Orlova, A., Agerer, B., Vilagos, B., Majek, P., Parapatics, K., Lercher, A. & Ringler, A. 2019. The ERBB-STAT3 axis drives Tasmanian devil facial tumor disease. Cancer Cell, 35, 125–139. e9.

Kowal, J., Arras, G., Colombo, M., Jouve, M., Morath, J. P., Primdal-Bengtson, B., Dingli, F., Loew, D., Tkach, M. & Thery, C. 2016a. Proteomic comparison defines novel markers to characterize heterogeneous populations of extracellular vesicle subtypes. Proc Natl Acad Sci U S A, 113, E968–77.

Kowal, J., Arras, G., Colombo, M., Jouve, M., Morath, J. P., Primdal-Bengtson, B., Dingli, F., Loew, D., Tkach, M. & Théry, C. 2016b. Proteomic comparison defines novel markers to characterize heterogeneous populations of extracellular vesicle subtypes. Proceedings of the National Academy of Sciences, 113, E968–E977.

Kubo, M., Kikuchi, K., Nashiro, K., Kakinuma, T., Hayashi, N., Nanko, H. & Tamaki, K. 1998. Expression of fibrogenic cytokines in desmoplastic malignant melanoma. The British journal of dermatology, 139, 192–197.

Kuroda, K., Okumura, K., Isogai, H. & Isogai, E. 2015. The human cathelicidin antimicrobial peptide LL-37 and mimics are potential anticancer drugs. Frontiers in oncology, 5, 144.

Kwon, Y. M., Stammnitz, M. R., Wang, J., Swift, K., Knowles, G. W., Pye, R. J., Kreiss, A., Peck, S., Fox, S., Pemberton, D., Jones, M. E., Hamede, R. & Murchison, E. P. 2018. Tasman-PCR: a genetic diagnostic assay for Tasmanian devil facial tumour diseases. R Soc Open Sci, 5, 180870.

Lazzaro, B. P., Zasloff, M. & Rolff, J. 2020. Antimicrobial peptides: Application informed by evolution. Science, 368.

Maas, S. L., Breakefield, X. O. & Weaver, A. M. 2017. Extracellular vesicles: unique intercellular delivery vehicles. Trends in cell biology, 27, 172–188.

Mccallum, H., Jones, M., Hawkins, C., Hamede, R., Lachish, S., Sinn, D. L., Beeton, N. & Lazenby, B. 2009. Transmission dynamics of Tasmanian devil facial tumor disease may lead to disease□induced extinction. Ecology, 90, 3379–3392.

Melo, S. A., Luecke, L. B., Kahlert, C., Fernandez, A. F., Gammon, S. T., Kaye, J., Lebleu, V. S., Mittendorf, E. A., Weitz, J. & Rahbari, N. 2015. Glypican-1 identifies cancer exosomes and detects early pancreatic cancer. Nature, 523, 177–182.

Minamida, S., Iwamura, M., Kodera, Y., Kawashima, Y., Ikeda, M., Okusa, H., Fujita, T., Maeda, T. & Baba, S. 2011. Profilin 1 overexpression in renal cell carcinoma. International Journal of Urology, 18, 63–71.

Montani, L., Buerki-Thurnherr, T., De Faria, J. P., Pereira, J. A., Dias, N. G., Fernandes, R., Gonçalves, A. F., Braun, A., Benninger, Y. & Böttcher, R. T. 2014. Profilin 1 is required for peripheral nervous system myelination. Development, 141, 1553–1561.

Murchison, E. P., Tovar, C., Hsu, A., Bender, H. S., Kheradpour, P., Rebbeck, C. A., Obendorf, D., Conlan, C., Bahlo, M., Blizzard, C. A., Pyecroft, S., Kreiss, A., Kellis, M., Stark, A., Harkins, T. T., Marshall Graves, J. A., Woods, G. M., Hannon, G. J. & Papenfuss, A. T. 2010. The Tasmanian devil transcriptome reveals Schwann cell origins of a clonally transmissible cancer. Science, 327, 84–7.

Norouzi-Barough, L., Asgari Khosro Shahi, A., Mohebzadeh, F., Masoumi, L., Haddadi, M. R. & Shirian, S. 2020. Early diagnosis of breast and ovarian cancers by body fluids circulating tumor-derived exosomes. Cancer Cell International, 20, 1–10.

Okumura, K., Itoh, A., Isogai, E., Hirose, K., Hosokawa, Y., Abiko, Y., Shibata, T., Hirata, M. & Isogai, H. 2004. C-terminal domain of human CAP18 antimicrobial peptide induces apoptosis in oral squamous cell carcinoma SAS-H1 cells. Cancer letters, 212, 185–194.

Ostrander, E. A., Davis, B. W. & Ostrander, G. K. 2016. Transmissible Tumors: Breaking the Cancer Paradigm. Trends in Genetics, 32, 1–15.

Pan, L.-H., Beppu, T., Kurose, A., Yamauchi, K., Sugawara, A., Suzuki, M., Ogawa, A. & Sawai, T. 2002. Neoplastic cells and proliferating endothelial cells express connective tissue growth factor (CTGF) in glioblastoma. Neurological research, 24, 677–683.

Peel, E., Cheng, Y., Djordjevic, J., Fox, S., Sorrell, T. & Belov, K. 2016. Cathelicidins in the Tasmanian devil (Sarcophilus harrisii). Scientific reports, 6, 35019.

Peinado, H., Aleckovic, M., Lavotshkin, S., Matei, I., Costa-Silva, B., Moreno-Bueno, G., Hergueta-Redondo, M., Williams, C., Garcia-Santos, G., Ghajar, C. M., Nitadori-Hoshino, A., Hoffman, C., Badal, K., Garcia, B. A., Callahan, M. K., Yuan, J. D., Martins, V. R., Skog, J., Kaplan, R. N., Brady, M. S., Wolchok, J. D., Chapman, P. B., Kang, Y. B., Bromberg, J. & Lyden, D. 2012. Melanoma exosomes educate bone marrow progenitor cells toward a pro metastatic phenotype through MET. Nature Medicine, 18, 883–+.

Pye, R. J., Pemberton, D., Tovar, C., Tubio, J. M., Dun, K. A., Fox, S., Darby, J., Hayes, D., Knowles, G. W., Kreiss, A., Siddle, H. V., Swift, K., Lyons, A. B., Murchison, E. P. & Woods, G. M. 2016. A second transmissible cancer in Tasmanian devils. Proc Natl Acad Sci U S A, 113, 374–9.

R CORE TEAM 2019. R: A language and environment for statistical computing. Vienna, Austria: R Foundation for Statistical Computing.

Rikkert, L. G., Nieuwland, R., Terstappen, L. & Coumans, F. 2019. Quality of extracellular vesicle images by transmission electron microscopy is operator and protocol dependent. Journal of extracellular vesicles, 8, 1555419.

Ruiz-Aravena, M., Jones, M. E., Carver, S., Estay, S., Espejo, C., Storfer, A. & Hamede, R. K. 2018. Sex bias in ability to cope with cancer: Tasmanian devils and facial tumour disease. Proceedings of the Royal Society B, 285, 20182239.

Safari, S., Baratloo, A., Elfil, M. & Negida, A. 2016. Evidence based emergency medicine; part 5 receiver operating curve and area under the curve. Emergency, 4, 111.

SAVE THE TASMANIAN DEVIL PROGRAM 2017. Risk Categorisation Guidelines for the keeping and movement of captive Tasmanian devils. 5.1 ed.: Department of Primary Industries, Park, Water & Environment.

Senol, O., Schaaij-Visser, T., Erkan, E., Dorfer, C., Lewandrowski, G., Pham, T., Piersma, S., Peerdeman, S., Ströbel, T. & Tannous, B. 2015. miR-200a-mediated suppression of non-muscle heavy chain IIb inhibits meningioma cell migration and tumor growth in vivo. Oncogene, 34, 1790–1798.

Silva, O. N., Porto, W. F., Ribeiro, S. M., Batista, I. & Franco, O. L. 2018. Host-defense peptides and their potential use as biomarkers in human diseases. Drug discovery today, 23, 1666–1671.

Takov, K., Yellon, D. M. & Davidson, S. M. 2019. Comparison of small extracellular vesicles isolated from plasma by ultracentrifugation or size-exclusion chromatography: yield, purity and functional potential. Journal of extracellular vesicles, 8, 1560809.

Thalmann, S., Peck, S., Wise, P., Potts, J. M., Clarke, J. & Richley, J. 2016. Translocation of a top-order carnivore: tracking the initial survival, spatial movement, homerange establishment and habitat use of Tasmanian devils on Maria Island. Australian Mammalogy, 38, 68–79.

Théry, C., Witwer, K. W., Aikawa, E., Alcaraz, M. J., Anderson, J. D., Andriantsitohaina, R., Antoniou, A., Arab, T., Archer, F. & Atkin-Smith, G. K. 2018. Minimal information for studies of extracellular vesicles 2018 (MISEV2018): a position statement of the International Society for Extracellular Vesicles and update of the MISEV2014 guidelines. Journal of extracellular vesicles, 7, 1535750.

Thery, C., Witwer, K. W., Aikawa, E., Alcaraz, M. J., Anderson, J. D., Andriantsitohaina, R., Antoniou, A., Arab, T., Archer, F., Atkin-Smith, G. K., Ayre, D. C., Bach, J. M., Bachurski, D., Baharvand, H., Balaj, L., Baldacchino, S., Bauer, N. N., Baxter, A. A., Bebawy, M., Beckham, C., Zavec, A. B., Benmoussa, A., Berardi, A. C., Bergese, P., Bielska, E., Blenkiron, C., Bobis-Wozowicz, S., Boilard, E., Boireau, W., Bongiovanni, A., Borras, F. E., Bosch, S., Boulanger, C. M., Breakefield, X., Breglio, A. M., Brennan, M. A., Brigstock, D. R., Brisson, A., Broekman, M. L. D., Bromberg, J. F., Bryl-Gorecka, P., Buch, S., Buck, A. H., Burger, D., Busatto, S., Buschmann, D., Bussolati, B., Buzas, E. I., Byrd, J. B., Camussi, G., Carter, D. R. F., Caruso, S., Chamley, L. W., Chang, Y. T., Chen, C. C., Chen, S., Cheng, L., Chin, A. R., Clayton, A., Clerici, S. P., Cocks, A., Cocucci, E., Coffey, R. J., Cordeiro-Da-Silva, A., Couch, Y., Coumans, F. A. W., Coyle, B., Crescitelli, R., Criado, M. F., D’souza-Schorey, C., Das, S., Chaudhuri, A. D., De Candia, P., De Santana, E. F., De Wever, O., Del Portillo, H. A., Demaret, T., Deville, S., Devitt, A., Dhondt, B., Di Vizio, D., Dieterich, L. C., Dolo, V., Rubio, A. P. D., Dominici, M., Dourado, M. R., Driedonks, T. A. P., Duarte, F. V., Duncan, H. M., Eichenberger, R. M., Ekstrom, K., Andaloussi, S. E. L., Elie-Caille, C., Erdbrugger, U., Falcon-Perez, J. M., Fatima, F., Fish, J. E., Flores-Bellver, M., Forsonits, A., Frelet-Barrand, A., et al. 2018. Minimal information for studies of extracellular vesicles 2018 (MISEV2018): a position statement of the International Society for Extracellular Vesicles and update of the MISEV2014 guidelines. Journal of Extracellular Vesicles, 7, 1535750.

Tovar, C., Obendorf, D., Murchison, E. P., Papenfuss, A. T., Kreiss, A. & Woods, G. M. 2011. Tumor-specific diagnostic marker for transmissible facial tumors of Tasmanian devils: immunohistochemistry studies. Vet Pathol, 48, 1195–203.

Von Haussen, J., Koczulla, R., Shaykhiev, R., Herr, C., Pinkenburg, O., Reimer, D., Wiewrodt, R., Biesterfeld, S., Aigner, A. & Czubayko, F. 2008. The host defence peptide LL-37/hCAP-18 is a growth factor for lung cancer cells. Lung cancer, 59, 12–23.

Weber, G., Chamorro, C. I., Granath, F., Liljegren, A., Zreika, S., Saidak, Z., Sandstedt, B., Rotstein, S., Mentaverri, R. & Sánchez, F. 2009. Human antimicrobial protein hCAP18/LL-37 promotes a metastatic phenotype in breast cancer. Breast cancer research, 11, 1–13.

Wenger, C., Ellenrieder, V., Alber, B., Lacher, U., Menke, A., Hameister, H., Wilda, M., Iwamura, T., Beger, H. G. & Adler, G. 1999. Expression and differential regulation of connective tissue growth factor in pancreatic cancer cells. Oncogene, 18, 1073–1080.

Wheeler, D. L., Barrett, T., Benson, D. A., Bryant, S. H., Canese, K., Chetvernin, V., Church, D. M., Dicuccio, M., Edgar, R. & Federhen, S. 2007. Database resources of the national center for biotechnology information. Nucleic acids research, 36, D13–D21.

Willms, E., Cabanas, C., Mager, I., Wood, M. J. A. & Vader, P. 2018. Extracellular Vesicle Heterogeneity: Subpopulations, Isolation Techniques, and Diverse Functions in Cancer Progression. Front Immunol, 9, 738.

Zanetti, M. 2005. The role of cathelicidins in the innate host defenses of mammals. Current issues in molecular biology, 7, 179–196.

Zasloff, M. 2002. Antimicrobial peptides of multicellular organisms. nature, 415, 389–395.

Zhou, B., Xu, K., Zheng, X., Chen, T., Wang, J., Song, Y., Shao, Y. & Zheng, S. 2020. Application of exosomes as liquid biopsy in clinical diagnosis. Signal transduction and targeted therapy, 5, 1–14.

Zhu, J., Chen, X., Liao, Z., He, C. & Hu, X. 2015. TGFBI protein high expression predicts poor prognosis in colorectal cancer patients. International Journal of Clinical and Experimental Pathology, 8, 702.

Zoidakis, J., Makridakis, M., Zerefos, P. G., Bitsika, V., Esteban, S., Frantzi, M., Stravodimos, K., Anagnou, N. P., Roubelakis, M. G. & Sanchez-Carbayo, M. 2012. Profilin 1 is a potential biomarker for bladder cancer aggressiveness. Molecular & cellular proteomics, 11, M111. 009449.

