## Supplementary Figures for "Cathelicidin-3 associated with serum extracellular vesicles enables early diagnosis of a transmissible cancer"

This file contains Supplementary figures 1–4

**Supplementary figure 1.**

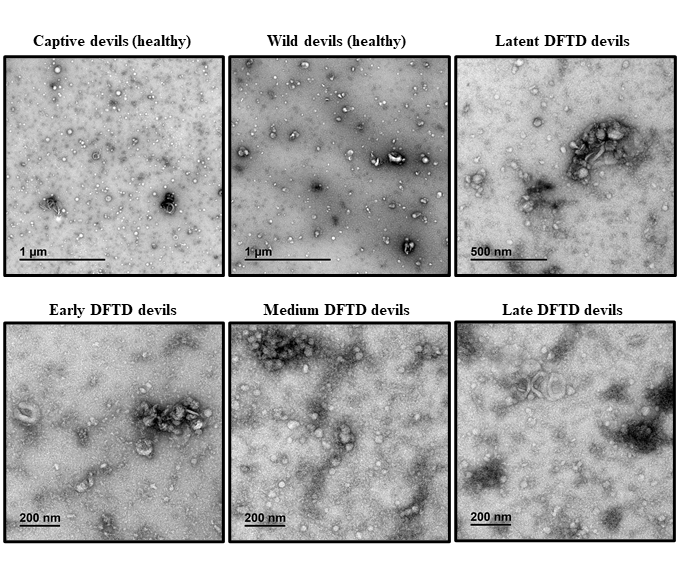
**Electron microscopy images of EVs derived from devil serum.** Transmission electron microscopy (TEM) images for EVs isolated from devil serum of healthy devils (captive and wild), and DFTD infected devils in early, medium, late, and latent stages of the disease. Four EV samples from different individuals were pooled for each group.

**Supplementary figure 2.**

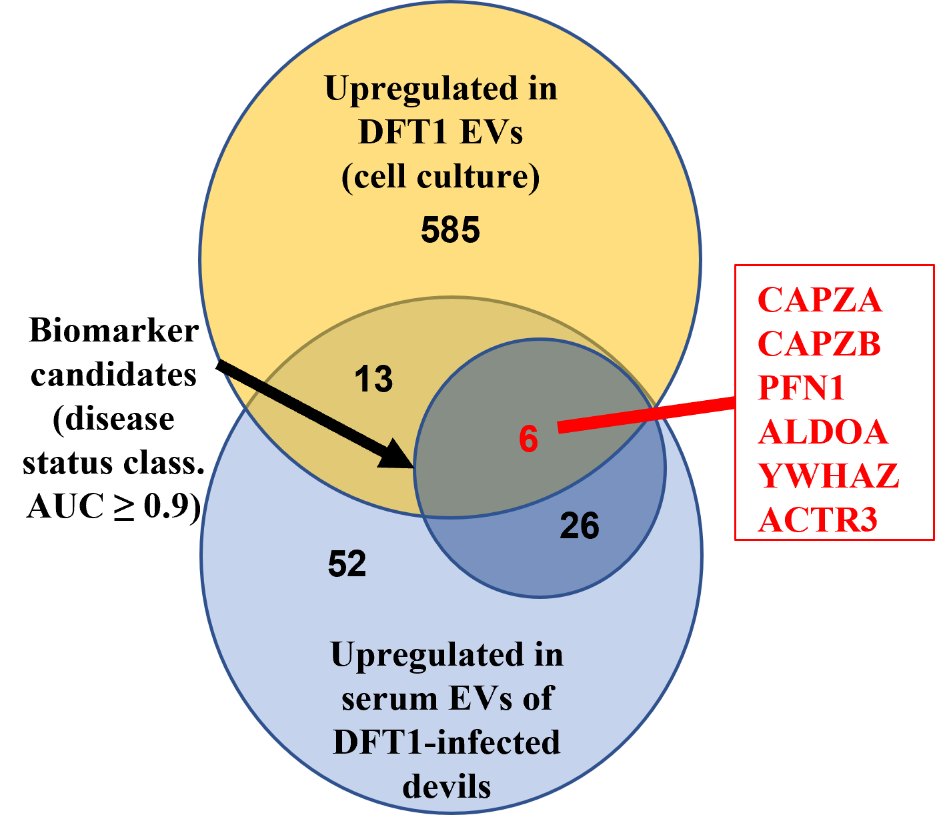

**Overlapping proteins from *in vitro* (DFT1 cell culture) and *in vivo* (devil serum) EV samples.** Venn diagram of overlapping proteins identified as upregulated in EVs derived from devil facial tumour cells relative to fibroblast cells and upregulated proteins in EVs derived from DFTD infected devils compared to captive obtained in a discovery cohort. Six of these 19 proteins yielded an area under the receiver operating characteristic curve greater or equal than 0.9.

**Supplementary figure 3.**

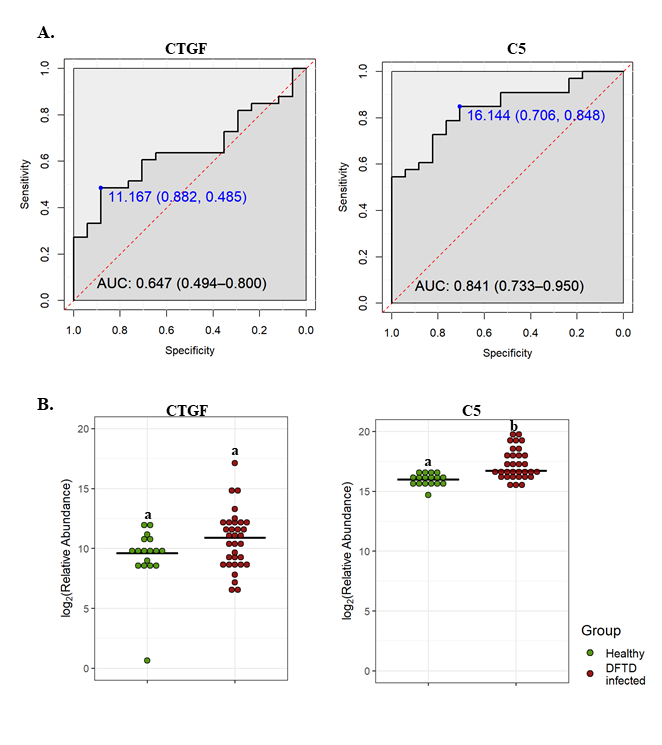

**CTGF and C5 showed a reduced performance in distinguishing different stages of DFT1. A.** Receiver operating characteristic (ROC) curve analysis for the EV proteins CTGF and C5 (DFTD vs healthy controls from the validation dataset). The dashed red identity line indicates random performance. The cut-off values were determined using Youden’s index and are indicated in blue at the left top corner of the ROC curve and specificity and sensitivity are indicated in brackets, respectively. **B.** Dot plots depicting the relative abundance of the EV proteins CTGF and C5 obtained from healthy animals and devils in different stages of DFTD (validation dataset), different letters “a” and “b” indicate significant differences between groups (Student’s t test, FDR-corrected p < 0.05).

**Supplementary figure 4.**

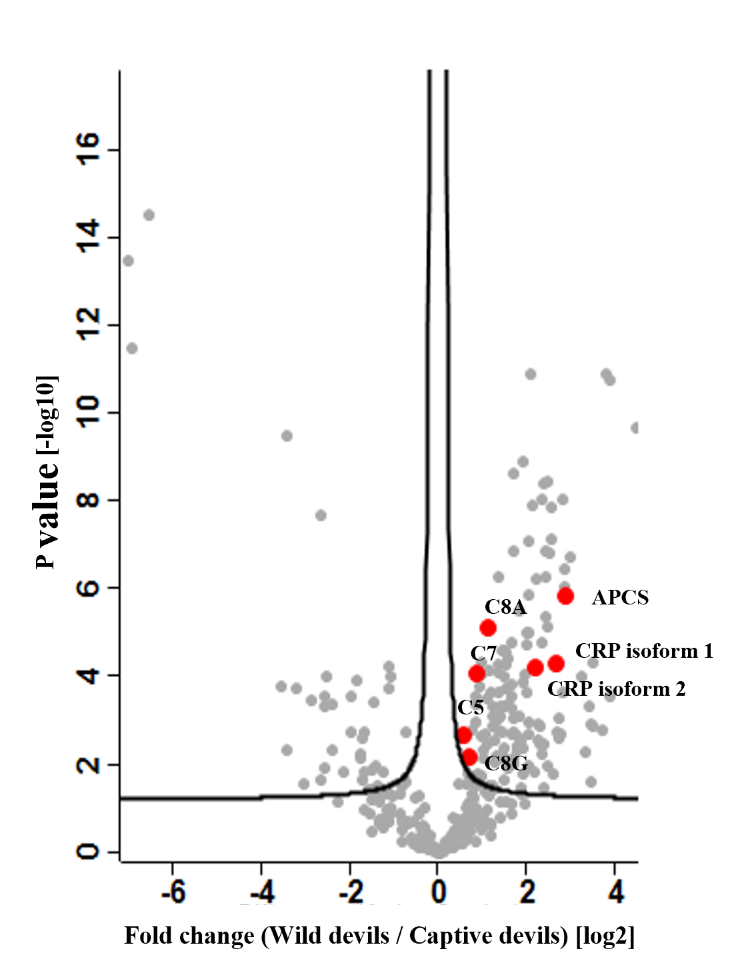

**Volcano plot of relative protein abundance in serum derived EVs of healthy captive relative to healthy wild devils.** Volcano plot of relative protein abundance in serum-derived EVs expressed as fold changes (log_2_) between healthy captive devils (n=10) and healthy wild devils (n=17) vs significance expressed as -log10 p-value. Proteins denoted in red are common inflammatory markers. APCS: serum amyloid P-component; CRP: C-reactive protein; C8A: complement C8 alpha chain, C8G: complement C8 gamma chain; C7: complement 7; C5: complement 5.
