## Supplementary Tables 2-4 for "Cathelicidin-3 associated with serum extracellular vesicles enables early diagnosis of a transmissible cancer"

**Supplementary Figures** for *Cathelicidin-3 associated with serum extracellular vesicles enables early diagnosis of a transmissible cancer***.**

This file contains Supplementary tables 2–4

**Supplementary table 2. EV proteins accuracy for the diagnosis of DFT1 in the discovery cohort with an area under the curve (AUC) ≥ 0.9**. All numbers for AUC, sensitivity, specificity, and accuracy are in fractions. 95% confidence intervals.

| Protein name | AUC | Specificity | Sensitivity | Accuracy | UniProt  number |
| --- | --- | --- | --- | --- | --- |
| CATH3 | 1.000 (1.000, 1.000) | 1.000 (1.000, 1.000) | 1.000 (1.000, 1.000) | 1.000 (1.000, 1.000) | G3W0S2 |
| CTGF | 1.000 (1.000, 1.000) | 1.000 (1.000, 1.000) | 1.000 (1.000, 1.000) | 1.000 (1.000, 1.000) | G3WBU2 |
| C5 | 1.000 (1.000, 1.000) | 1.000 (1.000, 1.000) | 1.000 (1.000, 1.000) | 1.000 (1.000, 1.000) | G3WZ02 |
| C8G | 0.992 (0.950, 1.000) | 1.000 (0.900, 1.000) | 0.917 (0.833, 1.000) | 0.955 (0.900, 1.000) | G3W1Q6 |
| SPP2 | 0.983 (0.925, 1.000) | 1.000 (0.900, 1.000) | 0.917 (0.833, 1.000) | 0.955 (0.900, 1.000) | G3VQP6 |
| C6 | 0.983 (0.925, 1.000) | 1.000 (0.800, 1.000) | 0.917 (0.833, 1.000) | 0.955 (0.909, 1.000) | G3WFL6 |
| C7 | 0.983 (0.925, 1.000) | 1.000 (0.800, 1.000) | 0.917 (0.833, 1.000) | 0.955 (0.909, 1.000) | G3WK50; G3WK51 |
| F10 | 0.983 (0.925, 1.000) | 1.000 (0.800, 1.000) | 0.917 (0.833, 1.000) | 0.955 (0.908, 1.000) | G3WXN7 |
| C8A | 0.975 (0.900, 1.000) | 1.000 (0.900, 1.000) | 0.917 (0.750, 1.000) | 0.955 (0.864, 1.000) | G3VVQ8; G3VVQ9 |
| ALDOA | 0.975 (0.900, 1.000) | 1.00 (0.800, 1.000) | 0.833 (0.750, 1.000) | 0.900 (0.864, 1.000) | G3WAK9 |
| C9 | 0.967 (0.900, 1.000) | 1.000 (0.700, 1.000) | 0.833 (0.750, 1.000) | 0.909 (0.864, 1.000) | G3W3E7 |
| ACTN1 | 0.958 (0.867, 1.000) | 1.000 (0.800, 1.000) | 0.833 (0.750, 1.000) | 0.909 (0.810, 1.000) | G3WFU1; G3WFU2 |
| CPNE1 | 0.958 (0.867, 1.000) | 0.800 (0.700, 1.000) | 1.000 (0.750, 1.000) | 0.909 (0.818, 1.000) | G3X2P2 |
| PLEK | 0.950 (0.825, 1.000) | 1.000 (1.000, 1.000) | 0.917 (0.750, 1.000) | 0.955 (0.864, 1.000) | G3VFV4; G3VFV5 |
| VCP | 0.942 (0.817, 1.000) | 1.000 (0.900, 1.000) | 0.833 (0.667, 1.000) | 0.909 (0.818, 1.000) | G3WW22 |
| MHC-I^*^ | 0.933 (0.808, 1.000) | 0.800 (0.600, 1.00) | 0.917 (0.667, 1.000) | 0.864 (0.773, 1.000) | G3VS26 |
| BIN2 | 0.925 (0.783, 1.000) | 0.900 (0.800, 1.000) | 0.917 (0.667, 1.000) | 0.909 (0773, 1.000) | G3VUC7 |
| LOC100931899 | 0.925 (0.792, 1.000) | 0.900 (0.700, 1.000) | 0.833 (0.583, 1.000) | 0.864 (0.773, 1.000) | G3VBX5 |
| PFN1 | 0.925 (0.792, 1.000) | 0.900 (0.700, 1.000) | 0.917 (0.667, 1.000) | 0.909 (0.773, 1.000) | G3VU42 |
| YWHAZ | 0.925 (0.766, 1.000) | 0.900 (0.700, 1.000) | 1.000 (1.000, 1.000) | 0.955 (0.864, 1.000) | G3WHE0 |
| TGFBI | 0.917 (0.767, 1.000) | 1.000 (0.700, 1.000) | 0.750 (0.583, 1.000) | 0.864 (0.773, 1.000) | G3VMG6 |
| IGH^*^ | 0.917 (0.775, 1.000) | 0.800 (0.600, 1.000) | 0.917 (0.583, 1.000) | 0.864 (0.773, 1.000) | G3VVK2 |
| PTN | 0.908 (0.758, 1.000) | 1.000 (0.600, 1.000) | 0.750 (0.583, 1.000) | 0.864 (0.773, 1.000) | G3VRT4; G3VRT5 |
| ANXA11 | 0.908 (0.767, 1.000) | 0.800 (0.600, 1.000) | 0.917 (0.583, 1.000) | 0.864 (0.773, 1.000) | G3VJ37; G3VJ38 |
| CAPZB | 0.908 (0.750, 1.000) | 0.900 (0.700, 1.000) | 0.917 (0.667, 1.000) | 0.909 (0.773, 1.000) | G3VTU6 |
| LOC100924827 | 0.908 (0.767, 1.000) | 0.900 (0.600, 1.000) | 0.750 (0.583, 1.000) | 0.818 (0.773, 1.000) | G3VT38 |
| TLN1 | 0.908 (0.758, 1.000) | 0.900 (0.700, 1.000) | 0.833 (0.583, 1.000) | 0.864 (0.773, 1.000) | G3WN56 |
| CAPZA2 | 0.900 (0.750, 1.000) | 0.900 (0.600, 1.000) | 0.833 (0.583, 1.000) | 0.864 (0.773, 1.000) | G3WVH2; G3WVH3 |
| MPO | 0.900 (0.750, 1.000) | 1.000 (0.600, 1.000) | 0.750 (0.583, 1.000) | 0.864 (0.773, 1.000) | G3WE24 |
| ANXA4 | 0.908 (0.758, 1.000) | 1.000 (0.800, 1.000) | 0.750 (0.583, 1.000) | 0.864 (0.773, 1.000) | G3WNV1; G3WNV2 |
| ACTR3 | 0.900 (0.750, 1.000) | 0.700 (0.600, 1.000) | 1.000 (0.583, 1.000) | 0.864 (0.773, 1.000) | G3WIY5 |

*Proteins blasted against the Tasmanian devil reference genome (GCA_902635505.1 mSarHar1.11) using the online NCBI protein Basic Local Alignment Search Tool (BLAST).

**Supplementary table 3.** **EV proteins accuracy for the diagnosis of DFTD in the validation cohort with an area under the curve (AUC) ≥ 0.9.** All numbers for AUC, sensitivity, specificity, and accuracy are in fractions. 95% confidence intervals.

| Protein  name | AUC | Specificity | Sensitivity | Accuracy | UniProt  number |
| --- | --- | --- | --- | --- | --- |
| CATH3 | 0.925  (0.840, 0.989) | 0.941  (0.882, 1.000) | 0.879  (0.727, 0.97) | 0.900  (0.820, 0.980) | G3W0S2 |
| PFN1 | 0.918  (0.804, 0.995) | 0.882  (0.765, 1.000) | 0.909  (0.788, 1.000) | 0.900  (0.820, 0.98) | G3VU42 |
| ILK | 0.914  (0.822, 0.980) | 0.824  (0.647, 1.000) | 0.848  (0.697, 1.000) | 0.840  (0.760, 0.960) | G3W679 |
| LIMS1 | 0.909  (0.818, 0.973) | 0.765  (0.646, 1.000) | 0.909  (0.636, 1.000) | 0.860  (0.740, 0.960) | G3VJH5 |

**Supplementary table 4. Kendall correlation of proteins with tumour volumes.** DFTD infected animals from the discovery and validation dataset (n=45). P values were adjusted by the Benjamini-Hochberg method.

| **Protein name** | **Kendall's tau** | **Corrected P value** | **Uniprot number** |
| --- | --- | --- | --- |
| MYH10 | 0.42 | 0.001 | G3W344 |
| TGFBI | 0.41 | 0.001 | G3VMG6 |
| CTGF | 0.41 | 0.001 | G3WBU2 |
| GADPH | 0.36 | 0.004 | G3WQA5 |
| HSPA5 | 0.36 | 0.004 | G3W4H8 |
| ANXA4 | 0.36 | 0.004 | G3WNV1;G3WNV2 |
| MYL6 | 0.35 | 0.006 | G3VPK8 |
| LOC100924827 | 0.34 | 0.006 | G3VT38 |
| MYH9 | 0.34 | 0.006 | G3W8S4 |
| ITGA2B | 0.34 | 0.008 | G3WU27 |
| RAP1A | 0.33 | 0.008 | G3WME1 |
| TNN | 0.33 | 0.008 | G3WK59 |
| FGL1 | 0.33 | 0.008 | G3W2N2 |
| RHOA | 0.33 | 0.008 | G3WUV1 |
| PPIA | 0.33 | 0.008 | G3WXF8 |
| LOC105750812 | 0.32 | 0.010 | G3VPL9 |
| LOC100931899 | 0.31 | 0.012 | G3VBX5 |
| PTN | 0.31 | 0.012 | G3VRT4; G3VRT5 |
| MHC-I^*^ | 0.30 | 0.017 | G3VS26 |
| ACTB | 0.29 | 0.018 | G3WHH2; G3WP12 |
| TUBB2A | 0.29 | 0.019 | G3WMM3 |
| YWHAQ | 0.29 | 0.020 | G3VS43 |
| ANXA1 | 0.28 | 0.022 | G3W8D2 |
| C5 | 0.28 | 0.022 | G3WZ02 |
| LOC100932693 | -0.28 | 0.022 | G3WEF1 |
| DCN | 0.27 | 0.029 | G3VVA7 |
| CCDC3 | 0.27 | 0.030 | G3WLM3 |
| DSTN | 0.27 | 0.032 | G3W5M4 |
| LIMS1 | 0.27 | 0.032 | G3VJH5; G3VJH6 |
| SELP | 0.27 | 0.032 | G3W7A0 |
| CFL1 | 0.26 | 0.034 | G3VHS2 |
| CAPZA2 | 0.26 | 0.035 | G3WVH2; G3WVH3 |
| CAPZB | 0.26 | 0.037 | G3VTU6 |
| YWHAZ | 0.26 | 0.037 | G3WHE0 |
| ARHGDIB | 0.26 | 0.039 | G3WV18 |
